# HPV18 E7 inhibits LATS1 kinase and activates YAP1 by degrading PTPN14

**DOI:** 10.1101/2024.03.07.583953

**Authors:** William J. Blakely, Joshua Hatterschide, Elizabeth A. White

**Affiliations:** Department of Otorhinolaryngology: Head and Neck Surgery, University of Pennsylvania Perelman School of Medicine, Philadelphia, PA, USA; Department of Integrative Immunobiology, Duke University School of Medicine, Durham, NC, USA

**Keywords:** Human papillomavirus, differentiation, tumor suppressor, keratinocyte, YAP1, Hippo, PTPN14

## Abstract

High-risk human papillomavirus (HPV) oncoproteins inactivate cellular tumor suppressors to reprogram host cell signaling pathways. HPV E7 proteins bind and degrade the tumor suppressor PTPN14, thereby promoting the nuclear localization of the YAP1 oncoprotein and inhibiting keratinocyte differentiation. YAP1 is a transcriptional coactivator that drives epithelial cell stemness and self-renewal. YAP1 activity is inhibited by the highly conserved Hippo pathway, which is frequently inactivated in human cancers. MST1/2 and LATS1/2 kinases form the core of the Hippo kinase cascade. Active LATS1 kinase is phosphorylated on threonine 1079 and inhibits YAP1 by phosphorylating it on amino acids including serine 127. Here, we tested the effect of high-risk (carcinogenic) HPV18 E7 on Hippo pathway activity. We found that either PTPN14 knockout or PTPN14 degradation by HPV18 E7 decreased phosphorylation of LATS1 T1079 and YAP1 S127 in human keratinocytes and inhibited keratinocyte differentiation. Conversely, PTPN14-dependent differentiation required LATS kinases and certain PPxY motifs in PTPN14. Neither MST1/2 kinases nor the putative PTPN14 phosphatase active site were required for PTPN14 to promote differentiation. Taken together, these data support that PTPN14 inactivation or degradation of PTPN14 by HPV18 E7 reduce LATS1 activity, promoting active YAP1 and inhibiting keratinocyte differentiation.

**SIGNIFICANCE:** The Hippo kinase cascade inhibits YAP1, an oncoprotein and driver of cell stemness and self-renewal. There is mounting evidence that the Hippo pathway is targeted by tumor viruses including human papillomavirus. The high-risk HPV E7 oncoprotein promotes YAP1 nuclear localization and the carcinogenic activity of high-risk HPV E7 requires YAP1 activity. Blocking HPV E7-dependent YAP1 activation could inhibit HPV-mediated carcinogenesis, but the mechanism by which HPV E7 activates YAP1 has not been elucidated. Here we report that by degrading the tumor suppressor PTPN14, HPV18 E7 inhibits LATS1 kinase, reducing inhibitory phosphorylation on YAP1. These data support that an HPV oncoprotein can inhibit Hippo signaling to activate YAP1 and strengthen the link between PTPN14 and Hippo signaling in human epithelial cells.

## INTRODUCTION

Human papillomaviruses are double-stranded DNA viruses with a tropism for keratinocytes in stratified squamous epithelia (1). The human papillomavirus (HPV) E6 and E7 proteins target host cellular proteins to reprogram cell signaling pathways, thereby enabling HPV replication and persistence. A subset of HPVs are the ‘high-risk’ genotypes that cause many anogenital and oropharyngeal cancers (2). High-risk HPV E6 and E7 proteins have biological activities that are not shared among the E6 and E7 proteins encoded by other HPVs, and high-risk HPV E6 and E7 expression can drive the immortalization and oncogenic transformation of human keratinocytes (1, 3).

One well-documented activity of many high-risk and low-risk (non-oncogenic) HPV E7 proteins is to bind and inactivate the retinoblastoma tumor suppressor (RB1) (4). RB1 inactivation drives the transcription of E2F target genes, thereby promoting the cell cycle progression that is required for HPV DNA replication (5, 6). In addition to binding RB1, high-risk HPV E7 proteins also target it for proteasome-mediated degradation (7). However, evidence from keratinocyte immortalization assays, other cell-based assays of HPV E7 transforming activity, and from studies of HPV E7 activity in transgenic mouse models supports that RB1 inactivation is necessary, but insufficient for the carcinogenic activity of high-risk HPV E7 (8–17).

We and others have determined that many HPV E7 proteins also target a second tumor suppressor, Protein Tyrosine Phosphatase Non-receptor Type 14 (PTPN14) (18–20). Several amino acids, including a highly conserved arginine in the HPV E7 C-terminus, enable E7 to bind directly to PTPN14 (21, 22). A conserved acidic amino acid in the HPV E7 N-terminus binds to the ubiquitin ligase UBR4, which is required for PTPN14 degradation and for the transforming activity of high-risk HPV E7 (13, 18). PTPN14 degradation by high-risk HPV E7 inhibits keratinocyte differentiation and extends primary epithelial cell lifespan, contributing to the immortalization of keratinocytes in culture (18, 21).

PTPN14 is mutated or its expression is reduced in several cancer types, and data from human cancer studies and cell-based assays provide evidence for the tumor suppressive activity of PTPN14 (23–34). The phosphatase domain of PTPN14 likely lacks enzymatic function and the putative catalytic site does not appear important for tumor suppressive activity (29, 30, 35). The best-characterized mechanism by which PTPN14 acts as a tumor suppressor is by inactivating the YAP1 oncoprotein, a potent driver of cell stemness and self-renewal (29, 31–34, 36). YAP1 is a transcriptional coregulator that is active when localized to the nucleus and bound to transcription factors including Transcriptional Enhanced Associate Domain (TEAD) proteins (37).

YAP1 and its paralog TAZ are normally repressed by the Hippo signaling pathway and Hippo pathway components are frequently mutated in human cancers (36, 38). The MST1/2 and LATS1/2 kinases form the core of the conserved Hippo kinase cascade (39, 40). MST1/2 activate LATS kinases at sites including threonine 1079 on LATS1 (41). Active LATS1/2 then phosphorylate YAP1 at serine 127 (42–45). LATS-dependent phosphorylation causes YAP1 to interact with 14-3-3 proteins and be retained in the cytoplasm where it is transcriptionally inactive and targeted for degradation (36, 38, 40, 45). Many additional protein-protein interactions regulate Hippo kinase activity and influence the assembly of Hippo components at the plasma membrane (36, 39). SAV1 promotes MST1 binding to LATS1 and MOB1A/B promote LATS1 binding to YAP1 (41, 46–49).

Other proteins including NF2 and WWC1/2/3 likely participate in and influence Hippo protein interactions (50, 51). The NF2 protein localizes to the plasma membrane in proximity to cytoskeletal scaffold elements and interacts with LATS1/2 (52–54). NF2 recruits LATS to the plasma membrane and promotes LATS kinase activity and Hippo signaling, but without affecting the MST kinase activity (51). WWC1 (KIBRA) and its paralogs WWC2 and WWC3 have been proposed to bind and regulate LATS kinase activity, perhaps by helping recruit LATS1/2 to the MST1/2-containing complex (55–57). Other data support that WWC proteins interact with NF2 (58–61). Of the three WWC paralogs, WWC1 has been studied most extensively (57).

Many of the interactions between Hippo components are mediated by binding of PPxY motifs in one protein to WW domains in another. Initial overexpression studies suggested that PPxY motifs in PTPN14 bound to WW domains in YAP1 (29, 31, 32). However, later biochemical experiments revealed that a separate set of PPxY motifs in PTPN14 make an exceptionally high-affinity interaction with WW motifs in WWC1, suggesting that the primary effect of PTPN14 does not occur through direct binding to YAP1 (62). The tumor suppressive activity of WWC1 requires PTPN14 (63).

Several observations highlight the potential role of YAP1, PTPN14, and the Hippo pathway in HPV carcinogenesis. We previously found that high-risk HPV E7 proteins promote the nuclear localization of YAP1 in the basal layer of stratified epithelial cell cultures, dependent on the ability of E7 to degrade PTPN14 (64). We also found that YAP1/TEAD-dependent transcriptional activity was required for high-risk HPV E7 proteins to extend the lifespan of primary human keratinocytes (64). Somatic mutation data in human cancers also argues for the influence of an HPV oncoprotein on Hippo signaling and YAP1. The three mitogenic pathways most differentially altered with high mutation rates in HPV-negative and low mutation rates in HPV-positive head and neck squamous cell carcinoma (HNSCC) are p53 (targeted by E6), cell cycle/RB1 (targeted by E7), and Hippo/YAP1 (64, 65).

Our findings support that RB1 inactivation and PTPN14 degradation are independent activities of high-risk HPV E7 that are each required for E7 transforming activity. The primary transcriptional readout in keratinocytes downstream of HPV E7-mediated PTPN14 degradation and YAP1 activation is a repression of epithelial differentiation. However, it was not known whether or how E7 and PTPN14 act on the Hippo pathway to regulate YAP1. Here we report that HPV18 E7 degrades PTPN14 to inhibit LATS1 kinase and activate YAP1, thereby inhibiting the commitment to keratinocyte differentiation. PTPN14 requires NF2 and its PY1/2 motif, which binds WWC1, to promote differentiation.

## RESULTS

### PTPN14 knockout reduces phosphorylation on YAP1 S127 and LATS1 T1079 in human keratinocytes

HPVs infect human keratinocytes and PTPN14 knockout or HPV E7-mediated PTPN14 degradation promotes YAP1 nuclear localization in the basal layer of stratified epithelia (64). However, the role of PTPN14 in Hippo signaling has not been tested in human keratinocytes, nor has the mechanism by which PTPN14 controls YAP1 been established. To test the hypothesis that PTPN14 degradation by E7 inhibits Hippo signaling, we monitored Hippo pathway activity in the presence and absence of PTPN14. First, we directly manipulated PTPN14. We transduced *hTERT* immortalized human keratinocytes (N/Tert-1) with lentiviral vectors to deliver Cas9 and one of two different sgRNAs targeting PTPN14, or non-targeting control sgRNAs. Disruption of the actin cytoskeleton activates Hippo kinases and we treated cells with the actin polymerization inhibitor cytochalasin D to induce Hippo signaling. YAP1 S127 phosphorylation (YAP1 pS127), which reflects decreased YAP1 activity, increased relative to total YAP1 during 90 minutes of cytochalasin D treatment in each nontargeting control cell line (Figure 1A). In contrast, YAP1 pS127 levels were lower in PTPN14 knockout cell lines treated with cytochalasin D. After 90 minutes of cytochalasin D treatment, there was a statistically significant decrease in the ratio of YAP1 pS127/total YAP1 in PTPN14 knockout cells compared to control cells (Figure 1B, Supplemental Table 1).

**Figure 1.**
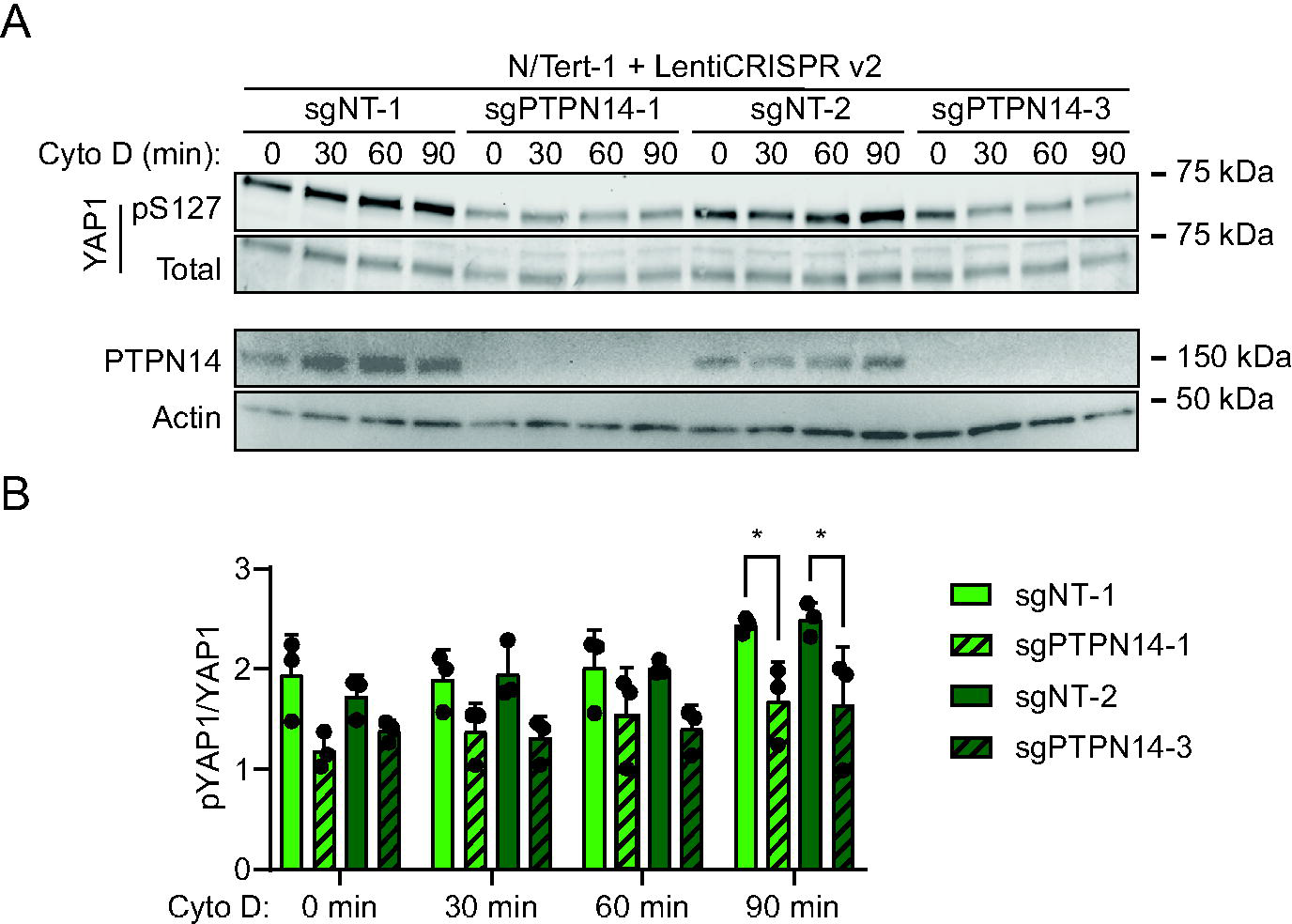
Phosphorylation on YAP1 S127 is reduced in PTPN14 knockout keratinocytes. N/Tert-1 keratinocytes were transduced with LentiCRISPRv2 vectors encoding spCas9 and an sgRNA sequence targeting PTPN14 or a nontargeting control sequence. sgRNA sequences were chosen from the Broad Institute Brunello Library. Four cell lines were generated using two different non-targeting control (sgNT-1, sgNT-2) and two different PTPN14 (sgPTPN14-1, sgPTPN14-3) sgRNAs. Cells were treated with cytochalasin D for up to 90 minutes and whole cell lysates were harvested at 30-minute intervals. (A) Whole cell protein lysates were separated by SDS-PAGE and proteins were detected by immunoblotting for YAP1, YAP1 pS127, PTPN14, and actin. (B) Western blot band intensity for three independent experiments was measured using ImageJ. Graph displays the ratio of YAP1 pS127/total YAP1 band intensity from three biological replicate experiments, plotted as mean ± standard deviation. Significance was determined by 2-way ANOVA with Holm-Šidak’s multiple comparisons test (*, P < 0.05)

Having observed that YAP1 pS127 was inhibited in PTPN14 knockout keratinocytes, we sought to determine whether PTPN14 knockout reduced phosphorylation on LATS, which is well-established as a marker of LATS kinase activity. To complement the cytochalasin D treatment experiments, we induced Hippo kinase activity by detachment of cells from their growth substrate (66–68). Two independent sets of nontargeting control or PTPN14 knockout N/Tert-1 cells were cultured at low density, then detached by trypsinization and incubated in suspension for 10 minutes. YAP1 pS127 levels increased upon suspension in control cells, but the increase in YAP1 pS127 was low or negligible in PTPN14 knockout cells (Figure 2A, B).

**Figure 2.**
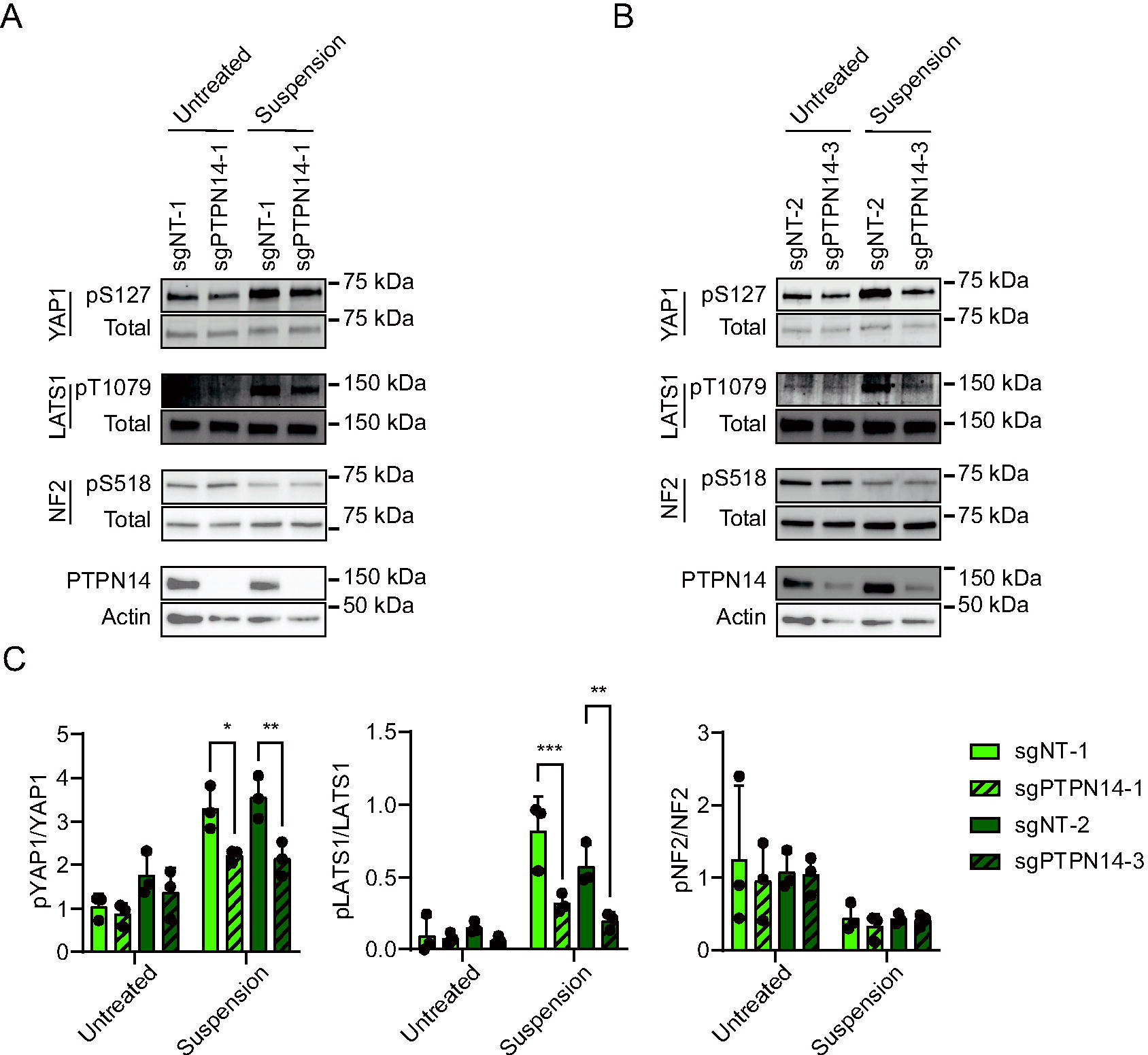
Phosphorylation on LATS1 T1079 is reduced in PTPN14 knockout keratinocytes. Control and PTPN14 knockout N/Tert-1 cells generated using LentiCRISPRv2 vectors were trypsinized and kept in suspension for 10 minutes. Whole cell protein lysates were separated by SDS-PAGE and proteins were detected by immunoblotting for YAP1, YAP1 pS127, LATS1, LATS1 pT1079, NF2, NF2 pS518, PTPN14, and actin. Panels (A, B) show replicate experiments using the same four distinct cell lines as in Figure 1. (C) Western blot band intensity for three independent experiments was measured using ImageJ. Graph displays the ratio of YAP1 pS127/total YAP1, LATS1 pT1079/total LATS1, or NF2 pS518/total NF2 band intensity from three biological replicate experiments, plotted as mean ± standard deviation. Significance was determined by 2-way ANOVA with Holm-Šidak’s multiple comparisons test (*, P < 0.05; **, P < 0.01; ***, P < 0.001)

Similarly, phosphorylation on LATS1 T1079 (LATS1 pT1079) showed a greater increase upon detachment in nontargeting control cells than in PTPN14 knockout cells. There was a statistically significant decrease in the ratio of YAP1 pS127/total YAP1 and in the ratio of LATS1 pT1079/total LATS1 in each PTPN14 knockout cell line compared to its matched control cell line (Figure 2C). LATS2 protein could not be detected by western blot in N/Tert-1 cells.

NF2 binds to LATS kinases and tethers them to the plasma membrane (53, 54, 57, 69).

Phosphorylation on NF2 S518 is reduced when Hippo signaling is active (70, 71). Dephosphorylated S518 indicates active NF2 and active Hippo signaling. Culturing cells in suspension decreased NF2 pS518 levels in both control and PTPN14 knockout N/Tert-1 cells (Figure 2), indicating that PTPN14 knockout alters phosphorylation on LATS1 and YAP1, but not NF2.

### HPV18 E7-mediated PTPN14 degradation reduced YAP1 S127 and LATS1 T1079 phosphorylation in human keratinocytes

To test whether PTPN14 degradation by a high-risk HPV E7 oncoprotein also reduces YAP1 and LATS1 phosphorylation, we used N/Tert-1 cells that express HPV18 E7, HPV18 E7 R84S, or a vector control. HPV18 E7 R84S cannot bind or degrade PTPN14 but can destabilize RB1 (21). Again, cytochalasin D treatment or culture in suspension were used to stimulate Hippo activity. YAP1 pS127 increased relative to total YAP1 over time in cytochalasin D-treated vector control cells, but not in wild type HPV18 E7 cells (Figure 3A, B). Both PTPN14 protein levels and the induction of YAP1 pS127 upon cytochalasin D treatment were partially restored in HPV18 E7 R84S cells. Stimulating Hippo kinase activity using cell detachment had a similar effect. YAP1 pS127 and LATS1 pT1079 increased after cell detachment in vector control and HPV18 E7 R84S cells, but the detachment-dependent increase was smaller in the wild type HPV18 E7 cells (Figure 3C, D). Overall, these data support that HPV18 E7 can inhibit Hippo-dependent phosphorylation on LATS1 T1079 and YAP1 S127, dependent on its ability to bind and degrade PTPN14.

**Figure 3.**
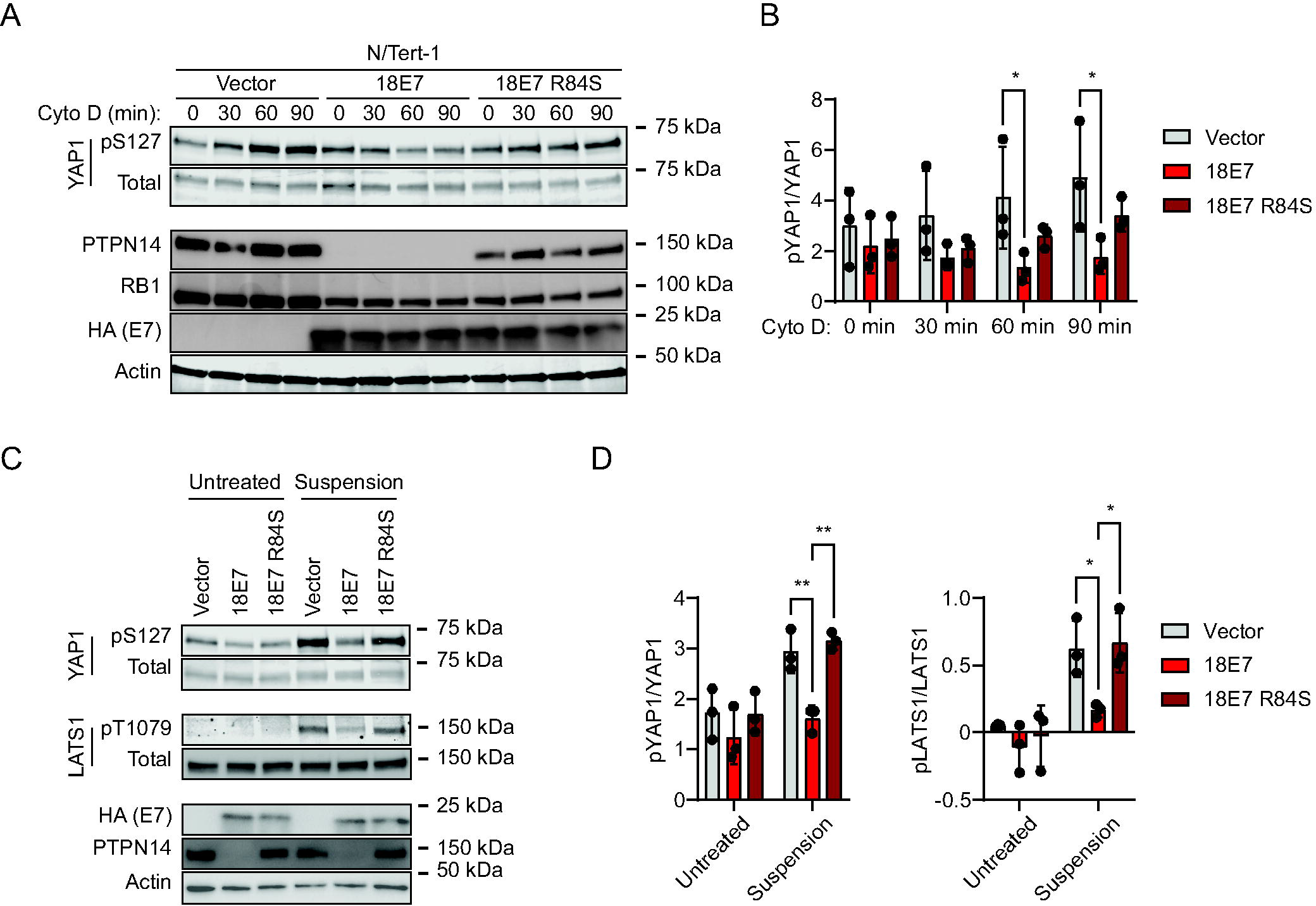
HPV18 E7-mediated PTPN14 degradation reduces phosphorylation on YAP1 S127 and LATS1 T1079. N/Tert-1 keratinocytes that stably express HA-tagged HPV18 E7 or HA-tagged HPV18 E7 R84S, which is unable to bind or degrade PTPN14, were used in assays of Hippo pathway activity. Cells transduced with empty vector were included as a control. (A) Cells were treated with cytochalasin D for up to 90 minutes and whole cell lysates were harvested at 30 minute intervals. Whole cell protein lysates were separated by SDS-PAGE and proteins were detected by immunoblotting for YAP1, YAP1 pS127, PTPN14, RB1, HA, and actin. (B) Western blot band intensity for three independent experiments was measured using ImageJ. Graph displays the ratio of YAP1 pS127/total YAP1 band intensity from three biological replicate cytochalasin D experiments, plotted as mean ± standard deviation. Significance was determined by 2-way ANOVA with Holm-Šidak’s multiple comparisons test (*, P < 0.05). (C) Cells were trypsinized and kept in suspension for 10 minutes. Whole cell protein lysates were separated by SDS-PAGE and proteins were detected by immunoblotting for YAP1, YAP1 pS127, LATS1, LATS1 pT1079, HA, PTPN14, and actin. (D) Western blot band intensity for three independent experiments was measured using ImageJ. Graph displays the ratio of YAP1 pS127/total YAP1 or LATS1 pT1079/total LATS1 band intensity from three biological replicate detachment experiments, plotted as mean ± standard deviation. Significance was determined by 2-way ANOVA with Holm-Šidak’s multiple comparisons test (*, P < 0.05; **, P < 0.01)

### PTPN14 promotes differentiation and Hippo pathway activity

PTPN14 knockout reduces the expression of epithelial differentiation genes and inhibits the expression of differentiation genes induced by cell detachment (18, 21). To test whether the effects of PTPN14 on differentiation correlate with its effects on YAP1 phosphorylation, we first tested the effect of PTPN14 knockout on differentiation in N/Tert-1 cells. Calcium treatment induces differentiation gene expression in keratinocytes and we confirmed that transcript levels for the differentiation marker genes Keratin 10 (*KRT10*) and involucrin (*IVL*) were reduced in calcium-treated PTPN14 knockout cells compared to matched controls (Figure 4).

**Figure 4.**
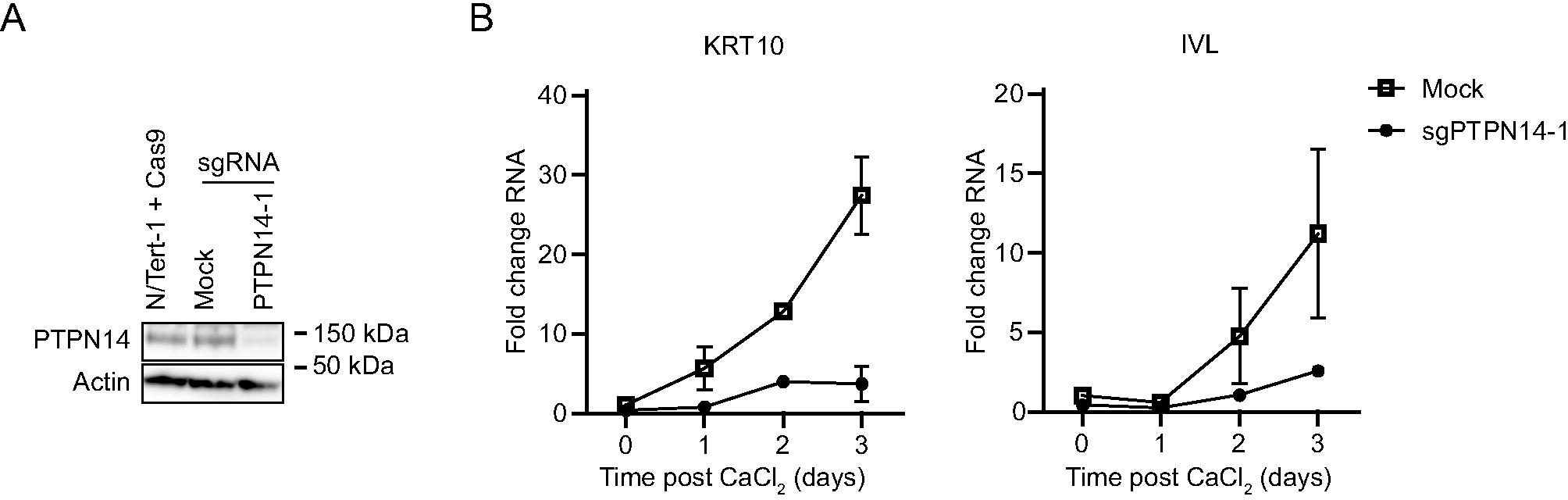
PTPN14 knockout impairs keratinocyte differentiation. (A) N/Tert-1 cells stably expressing Cas9 were transfected with sgRNA targeting PTPN14 (Synthego) or mock transfected and PTPN14 knockout was validated by immunoblot. (B) Mock transfected and PTPN14 knockout N/Tert-1 cells were cultured in media containing 1.5 mM calcium and harvested for RNA analysis at indicated times. KRT10 and IVL RNA levels were analyzed by qRT-PCR and normalized to G6PD. Graphs show values for technical duplicate experiments. Error bars display mean ± range.

The observation that PTPN14 limits the expression of differentiation genes prompted us to test whether the converse was also true: that elevated PTPN14 levels could increase differentiation gene expression. N/Tert-1 keratinocytes were transduced with a lentivirus encoding HA-tagged doxycycline-inducible PTPN14 (pLIX-PTPN14). *IVL* and *KRT10* transcript levels increased in doxycycline-treated N/Tert-pLIX-PTPN14 cells (Figure 5A), a finding consistent with our previous report that PTPN14 overexpression increased the expression of differentiation marker genes *KRT1* and *IVL* in primary human foreskin keratinocytes (HFK) (64). Having established that increased PTPN14 could promote differentiation in N/tert-1 cells, we tested whether it altered phosphorylation on Hippo proteins. Doxycycline treatment induced PTPN14, increased YAP1 pS127 and LATS1 pT1079, and modestly decreased NF2 pS518 (Figure 5B). Overall, PTPN14 overexpression increased phosphorylation on YAP1 pS127 and LATS1 pT1079 and promoted differentiation gene expression. Conversely, PTPN14 knockout inhibited YAP1 pS127 and LATS1 pT1079 and reduced differentiation gene expression.

**Figure 5.**
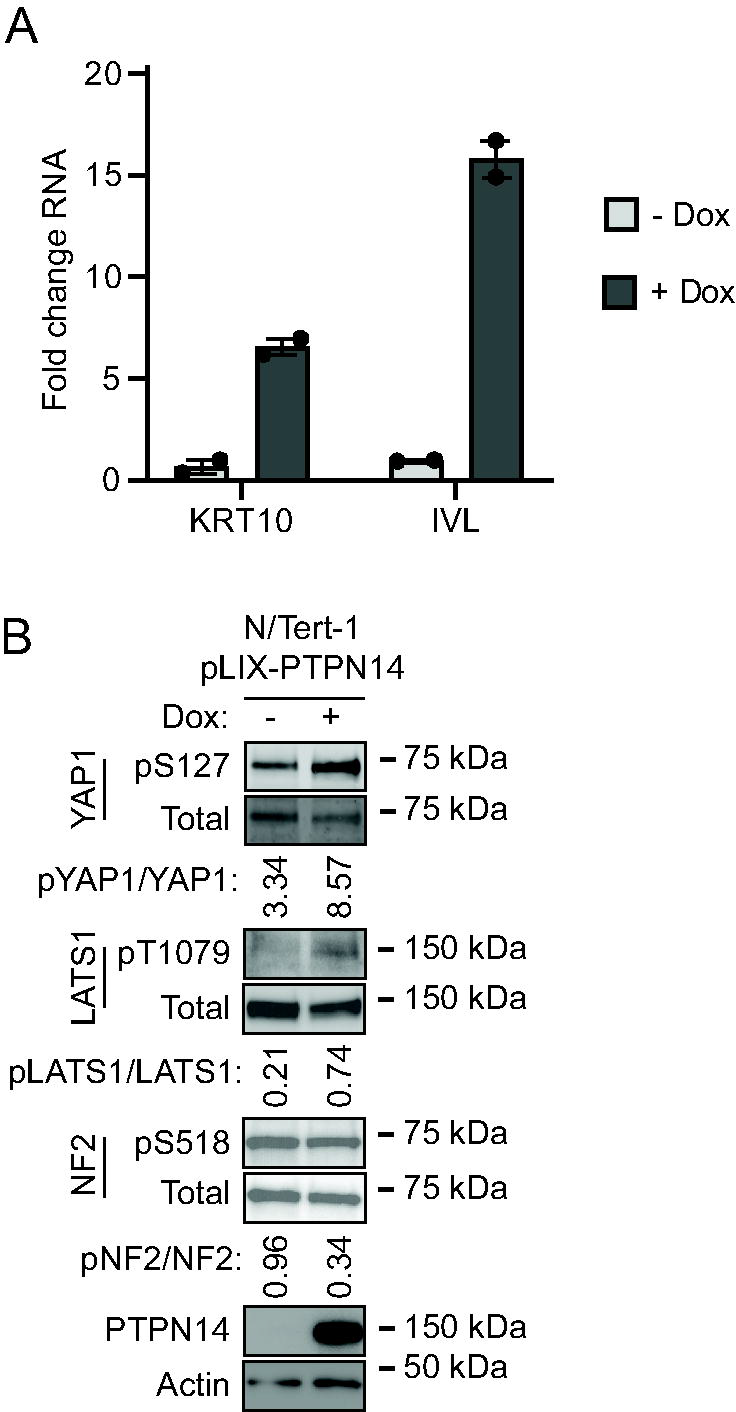
PTPN14 promotes keratinocyte differentiation and Hippo pathway activity. N/Tert-1 keratinocytes that express endogenous PTPN14 were transduced with a lentiviral vector encoding doxycycline-inducible PTPN14 (pLIX-PTPN14). Cells were treated with 1 µg/mL doxycycline for 24 hours or left untreated. (A) KRT10 and IVL RNA levels were measured by qRT-PCR and normalized to GAPDH. Graphs show data points for two technical replicate experiments. Error bars display mean ± range. (B) Whole cell protein lysates were separated by SDS-PAGE and proteins analyzed by immunoblotting for YAP1, YAP1 pS127, LATS1, LATS1 pT1079, NF2, NF2 pS518, PTPN14, and actin. Bands for YAP1, YAP1 pS127, LATS1, LATS1 pT1079, NF2, NF2 pS518 were quantified by densitometry. Values reflect the ratio of phospho protein/total protein band density.

### The PTPN14 PY1/2 motifs are required to induce keratinocyte differentiation

Next we used PTPN14 overexpression to determine which region(s) of PTPN14 upregulate differentiation. Several protein components of the Hippo pathway interact with one another via complementary WW domains and PPxY (PY) motifs (62, 72). PTPN14 contains two sets of PY motifs and has a C-terminal protein tyrosine phosphatase domain. However, the amino acid sequence of the PTPN14 phosphatase domain diverges from the consensus sequence in active protein tyrosine phosphatases and it is likely to have little or no catalytic activity (35). We designed two mutants, each lacking a PY motif pair, and a third containing a serine point mutant C1121S at the conserved reactive cysteine of the putative phosphatase active site (Figure 6A) (73, 74). We transduced HFKs with lentiviruses encoding wild-type (WT) and mutant PTPN14, then measured *KRT1* differentiation marker transcript levels (Figure 6B). *KRT1* levels increased in WT PTPN14, PTPN14 ΔPY3/4, and PTPN14 C1121S cells, but not in PTPN14 ΔPY1/2 cells.

**Figure 6.**
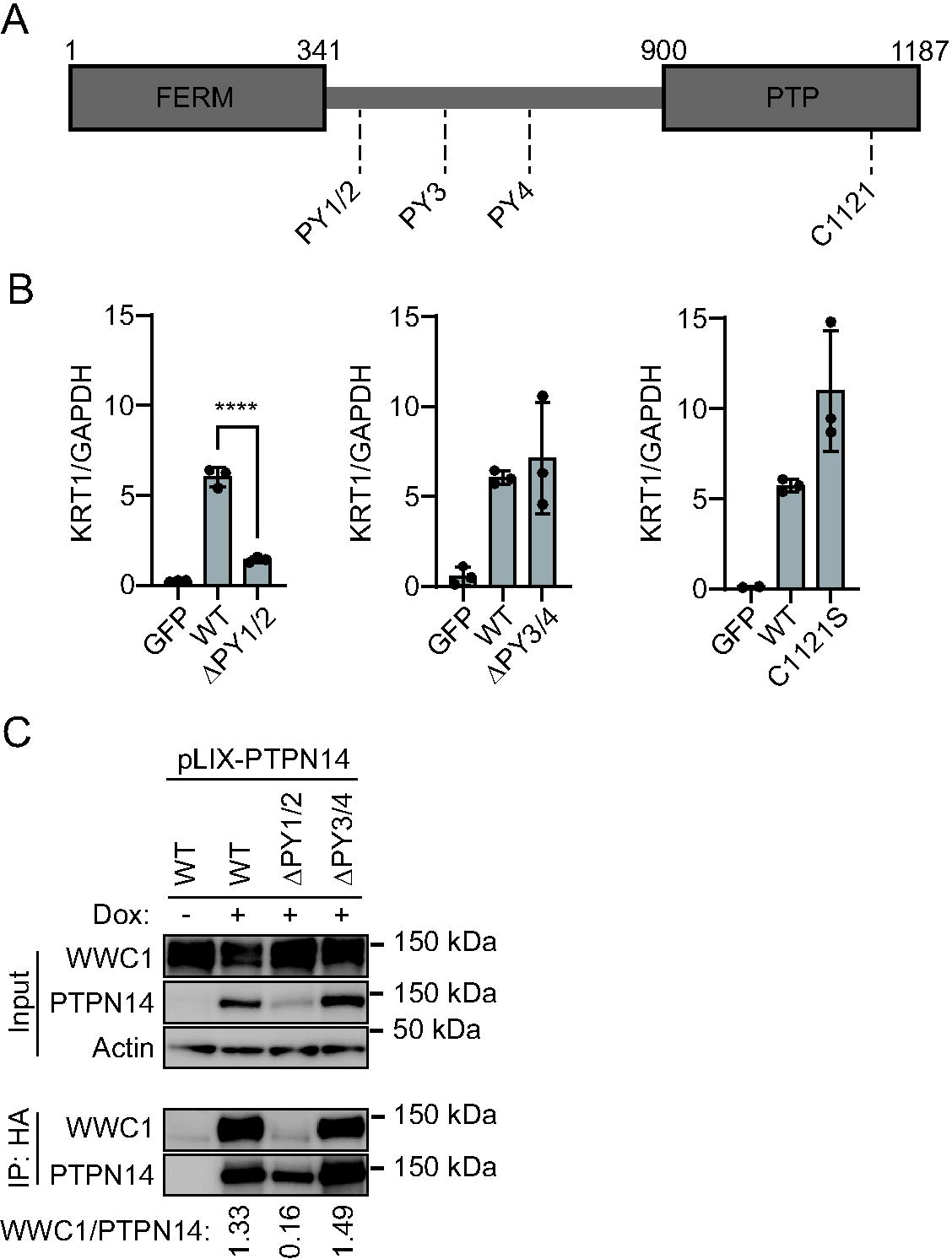
PTPN14 PY1/2 motifs are required to induce differentiation. (A) Schematic of PTPN14 domains and location of PPxY motifs. (B) HFK were transfected with lentiviral constructs encoding GFP, wild-type (WT) PTPN14, or PTPN14 mutants as indicated. Total cellular RNA was collected and KRT1 transcript levels were analyzed by qRT-PCR and normalized to GAPDH. Graph shows individual data points for three independent experiments. Data is plotted as mean ± standard deviation. Significance was determined by ANOVA with Holm-Šidak’s multiple comparisons test. (ns, not significant; ****, P < 0.0001). (C) N/Tert-1 immortalized keratinocytes transduced with pLIX-PTPN14 were treated with 1 µg/mL doxycycline for 48 hours to induce HA-tagged PTPN14 expression. Whole cell protein lysates were collected and PTPN14 immunoprecipitated with anti-HA agarose beads. Protein lysates from input and elution fractions were separated by SDS-PAGE and proteins were analyzed by immunoblotting for WWC1, PTPN14, and actin. Bands for WWC1 and PTPN14 in the IP fraction were quantified by densitometry. Values reflect the ratio of WWC1/PTPN14 band intensity.

The PTPN14 PY1/2 motifs bind with high affinity to the WW domain region of WWC1 (KIBRA) (62). We therefore tested whether WT or PY mutant forms of PTPN14 bound to WWC1 in human keratinocytes. N/Tert-Cas9 cells were transfected with a PTPN14 sgRNA to eliminate endogenous PTPN14, then transduced with pLIX vectors encoding WT or mutant HA-tagged PTPN14. Although the steady-state level of PTPN14 ΔPY1/2 was lower than that of WT PTPN14 or PTPN14 ΔPY3/4, some PTPN14 ΔPY1/2 was expressed and immunoprecipitated in the experiment. PTPN14 and KRT10 protein levels increased upon induction of WT PTPN14 (Figure S1). Cells were treated with doxycycline and lysates subjected to anti-HA immunoprecipitation. Wild-type PTPN14 and PTPN14 ΔPY3/4 co-immunoprecipitated comparable amounts of WWC1 (Figure 6C). Even accounting for the lower expression of the PTPN14 ΔPY1/2 mutant, a negligible amount of WWC1 was co-immunoprecipitated by PTPN14 PY1/2 compared to WT PTPN14 or PTPN14 ΔPY3/4. We conclude that the ability of PTPN14 to induce keratinocyte differentiation requires the PY motifs that bind to WWC1, but not the PTPN14 C1121 residue.

### LATS kinases are required for PTPN14 to induce keratinocyte differentiation

We previously used siRNA knockdown combined with PTPN14 overexpression to demonstrate that YAP1 is required for PTPN14 to promote differentiation gene expression (64). The finding was based on the observation that although knockdown of YAP1 and its paralog TAZ increased differentiation gene expression, PTPN14 overexpression could not further increase the levels of differentiation genes in YAP1/TAZ-depleted cells. Our data further supported that the predominant effect of PTPN14 on differentiation is mediated by YAP1, not TAZ. We also observed that knockdown of either LATS1 or LATS2 in HFK reduced basal levels of differentiation gene expression (64).

Here we used the same knockdown plus overexpression strategy to determine which of several additional components of the Hippo pathway are required for PTPN14 to induce keratinocyte differentiation. We transfected HFKs with siRNA targeting YAP1 and TAZ, LATS1 and LATS2, or a non-targeting control. Twenty-four hours post transfection, cells were transduced with lentiviruses encoding PTPN14 or GFP, then total cellular RNA was harvested 72h post transfection. In siControl treated cells, PTPN14 expression increased the levels of *KRT10*, *KRT1*, and *IVL* (Figure 7, Figure S2). Consistent with our previous finding, YAP1 and TAZ knockdown in HFK increased the expression of differentiation genes, but PTPN14 could not further increase differentiation in cells depleted of YAP1/TAZ compared to those transfected with siControl. Although PTPN14 retained some ability to increase *KRT10, KRT1, and IVL* RNA in cells treated with two different sets of siRNAs targeting LATS1 and LATS2 (Figure 7A, Figure S2), levels of all three differentiation marker genes remained low in LATS knockdown cells throughout the experiment. In the absence of LATS kinases, PTPN14 did not increase the levels of *KRT10, KRT1, or IVL* beyond the amount observed in untreated control cells. Knockdowns and PTPN14 overexpression were validated by qRT-PCR (Figure S3).

**Figure 7.**
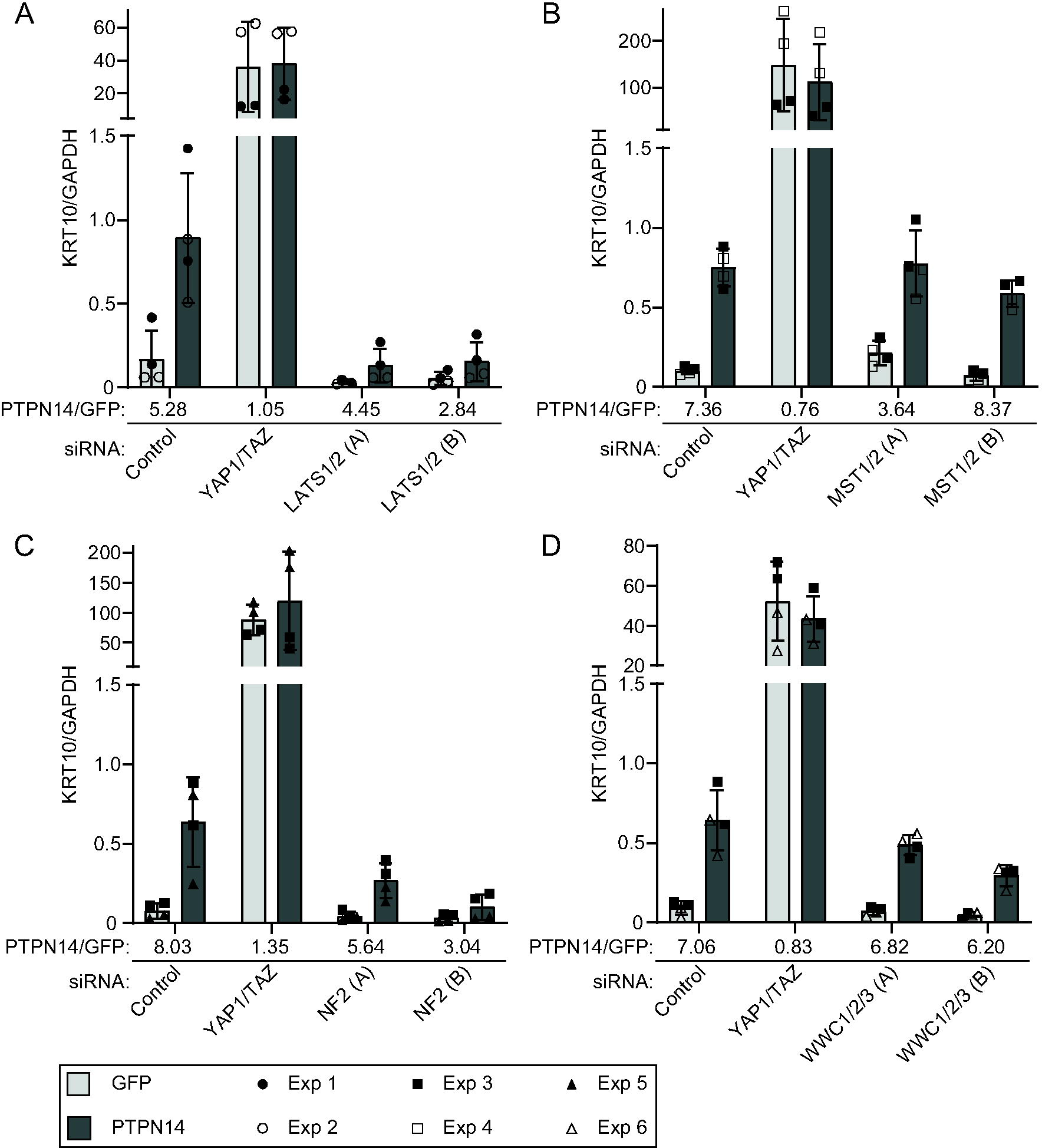
LATS kinases and NF2 are required for PTPN14 to induce KRT10 expression. HFK were transfected with siRNA then transduced with lentiviruses encoding GFP or PTPN14 at 24h post transfection. Total cellular RNA was collected 72h post-knockdown and 48h post-transduction. RNA transcripts for KRT10 were measured by qRT-PCR and normalized to GAPDH. Six individual experiments were conducted, each in technical duplicate. Each experiment included an siControl condition, a YAP1 and TAZ siRNA-treated condition, and siRNAs targeting additional component(s) of the Hippo pathway. Each component of the Hippo pathway was targeted with two different siRNAs per gene, denoted A and B. Panels display data from knockdowns as follows: (A) LATS1 and LATS2, (B) MST1 and MST2, (C) NF2, or (D) WWC1, WWC2, and WWC3. Data are graphed as mean ± standard deviation of combined replicate data. PTPN14/GFP denotes the ratio of KRT10 level in PTPN14 transduced cells vs GFP transduced cells in each siRNA-treated condition. Since Experiment 3 included several siRNAs (Control, YAP1/TAZ, MST1/2, NF2, WWC1/2/3), the same siControl and siYAP1/TAZ data from experiment 3 is included in panels B, C, and D.

Next we tested whether PTPN14 required MST1 and MST2, NF2, or WWC1 and its paralogs WWC2 and WWC3 to induce differentiation. We transfected HFK with siRNA targeting MST1 and MST2, NF2, or all three WWC genes, transduced the cells with lentiviruses encoding GFP or PTPN14, and measured *KRT10* transcript levels. Depletion of MST1 and 2 did not interfere with the ability of PTPN14 to induce *KRT10* (Figure 7B). Similar to the knockdown of LATS1 and LATS2, knockdown of NF2 limited, although did not completely eliminate, the ability of PTPN14 to induce *KRT10* (Figure 7C). Depletion of WWC genes had an intermediate effect, but PTPN14 was still able to induce *KRT10* several-fold in the absence of WWC1/2/3. The magnitude of reduction in PTPN14-induced differentiation correlated with the magnitude of depletion of WWC1 (Figure 7D, Figure S3). These data support that LATS kinases and NF2 are required for full induction of differentiation by PTPN14.

### PTPN14 knockout promotes anchorage independent growth and reduces YAP1 pS127 in HEK TER cells

Finally, we tested the effect of PTPN14 knockout in a human cell transformation assay. HEK TER cells are human embryonic kidney (HEK) cells that express the catalytic subunit of human telomerase (*hTERT*), Simian Virus 40 (SV40) Large T antigen (LT), and an oncogenic allele of *ras* (H-*ras* V12) (75, 76). The cells are immortalized but do not grow without attachment to a substrate. HEK TER cells acquire the capacity for anchorage-independent growth upon expression of SV40 Small T (ST) antigen. ST activates YAP1 and transforms HEK TER cells by interacting with a PP2A subunit of the STRIPAK complex upstream of the Hippo pathway (77). We previously reported that HPV16 E7 could promote the anchorage-independent growth of HEK TER cells (11). The HPV16 E7 transforming activity in HEK TER cells mapped broadly to the regions required to degrade PTPN14 (13, 19).

To test whether PTPN14 knockout promotes the anchorage-independent growth of HEK TER cells, we generated CRISPR/Cas9 edited HEK TER cells that do not express PTPN14.

LATS1/2 knockout HEK TER cells were included to verify that Hippo pathway inhibition promoted HEK TER growth and HEK TER cells stably expressing SV40 ST (11) were included as a positive control. HEK TER cells depleted of PTPN14, LATS1/2, or that expressed SV40 ST readily formed colonies in soft agar assays (Figure 8A), whereas HEK TER cells transfected with non-targeting sgRNA exhibited minimal anchorage-independent growth. Consistent with our findings in human keratinocytes and with the previous report that SV40 ST reduces YAP1 phosphorylation (77), cytochalasin D treatment induced YAP1 pS127 more strongly in HEK TER sgNT cells than in cells depleted of PTPN14, LATS1 and LATS2, or that expressed SV40 ST (Figure 8B, C). Using a second set of sgRNAs directed against PTPN14 and LATS1 and LATS2 resulted in a similar trend in colony formation (Figure S4).

**Figure 8.**
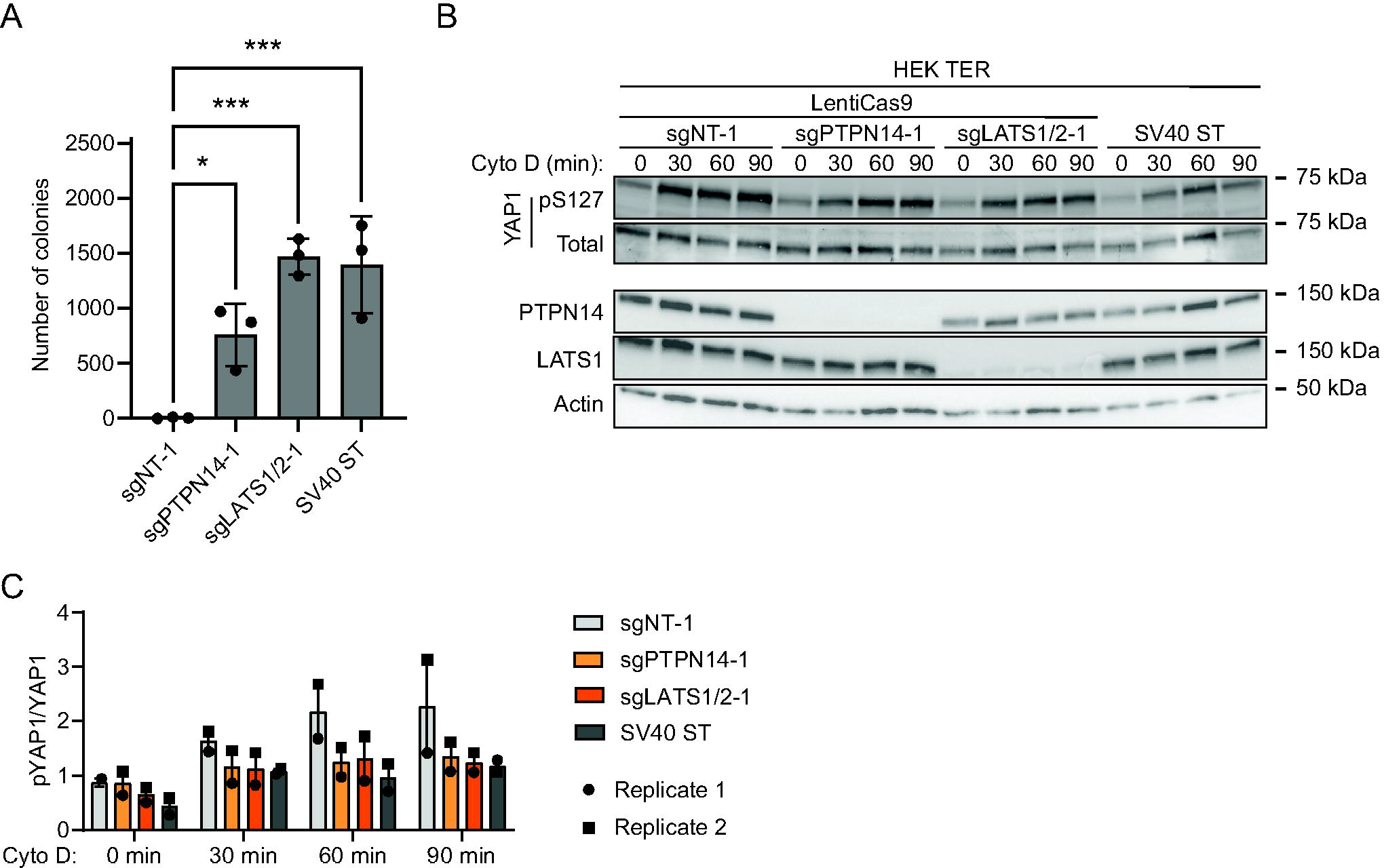
PTPN14 knockout promotes anchorage independent growth and reduces YAP1 phosphorylation in HEK TER cells. (A) HEK TER cells expressing spCas9 were transfected with sgRNA targeting PTPN14 or LATS1 and LATS2. HEK TER cells expressing SV40 ST were used as a positive control for colony formation. Cells were plated in soft agar in technical triplicate and incubated at 37°C for 18 days, then photographed. Colonies were counted and quantified using ImageJ software. Graphs show individual data points for each plate and indicate mean ± standard deviation. Statistical significance of the nontargeting control condition compared to experimental conditions was determined by ANOVA with Dunnett’s multiple comparisons test (*, P < 0.05; ***, P < 0.001). (B) Cells were treated with cytochalasin D for up to 90 minutes and whole cell lysates were harvested at 30-minute intervals. Whole cell protein lysates were separated by SDS-PAGE and proteins were detected by immunoblotting for YAP1, YAP1 pS127, PTPN14, LATS1, and actin. (C) Band intensity for blots of two independent experiments was measured by ImageJ. Graph displays the ratio of YAP1 pS127/total YAP1 band intensity from two biological replicate cytochalasin D experiments as mean ± range. The blots shown in panel B are from experiment 1.

## DISCUSSION

YAP1 is a transcriptional coactivator that promotes epithelial cell stemness and self-renewal, limiting differentiation (42, 50, 78–85). We previously reported that oncogenic high-risk HPV E7 proteins bind and target PTPN14 for proteasome-mediated degradation (19, 86). The HPV18 E7-mediated degradation of PTPN14 results in increased nuclear YAP1 localization and the maintenance of basal epithelial identity in three-dimensional models of stratified epithelial tissue (64). However, we had not tested whether or how HPV E7-mediated degradation of PTPN14 alters Hippo pathway activity, which controls YAP1, in keratinocytes.

Several reports indicate that PTPN14 can inhibit YAP1 activity, although the mechanistic basis of such inhibition is incompletely understood (24, 29–34, 55, 64). Initial studies suggested that PTPN14 bound directly to YAP1 via the second set of PTPN14 PY motifs (PY3/4 in this paper), but more recent analysis of PY-WW interactions among Hippo components determined that the PTPN14 PY1/2 tandem motif has nanomolar affinity for WW domains in Hippo components including WWC1 (31–33, 62). These and other data support that PTPN14, WWC1, and LATS1/2 work together to inhibit YAP1 (24, 55, 56), but not by binding YAP1 directly.

Certain assays of tumor suppression show a mutual requirement for PTPN14 and WWC1 (KIBRA) (55, 62, 63, 87, 88), but potential contributions of WWC2 and WWC3 in the same assays have not been thoroughly tested (59). Other data indicate that PTPN14 and/or WWC proteins act through NF2, which interacts with LATS1/2 and recruits LATS1/2 to a plasma membrane-associated complex that also contains MST1/2 and its cofactor SAV1 (51, 60, 61, 70, 89–91).

Our data strengthen the model that PTPN14 regulates LATS kinases in a manner that also requires NF2 and the PTPN14 PY1/2 motif, which binds WWC1. In support of this model, we found that either PTPN14 knockout or HPV18 E7-mediated PTPN14 degradation reduced the levels of LATS1 pT1079 and YAP1 pS127, indicating that PTPN14 degradation reduces Hippo activity and activates YAP1 (Figure 1, Figure 2, Figure 3). The effect of HPV18 E7 on YAP1 was independent of its ability to degrade RB1 (Figure 3). Conversely, PTPN14 overexpression increased phosphorylation on LATS1 T1079 and YAP1 S127 (Figure 5).

We established the connection between PTPN14, YAP1 activity, and keratinocyte differentiation (Figure 4, Figure 5, (18)) and used keratinocyte differentiation as a readout to probe the effect of PTPN14 on Hippo signaling. We found that the induction of differentiation by PTPN14 required its PY1/2 motifs, which bind WWC1, but not the catalytic cysteine of the putative active site (Figure 6). PTPN14 required LATS1/2 kinases and NF2 to promote differentiation (Figure 7), further supporting that PTPN14 acts on LATS kinases only in the presence of NF2. Our finding that PTPN14 could still induce differentiation in cells transfected with MST1/2 siRNAs is consistent with the finding that MST1/2 are not required for PTPN14 overexpression to induce LATS1 phosphorylation (55). It is possible that one of several other kinases can phosphorylate LATS in the absence of MST1/2 (59, 92–96). There at least two separate branches of the Hippo pathway that can activate LATS1/2, one converging on MST1/2 kinases and the other converging on MAP4K4 (95–98). MST1/2 kinases are not active in stratified epithelial cells (99, 100), and further experiments will be required to determine whether PTPN14 degradation alters the activity of the MAP4K4 branch of the Hippo pathway.

PTPN14 could still promote differentiation in cells treated with WWC1/2/3 siRNA, even though the WWC1 binding motif was required for PTPN14 to promote differentiation (Figure 6B, Figure 7). It is possible that incomplete siRNA-based depletion of WWC1/2/3 resulted in a weaker phenotype than that observed with the PTPN14 ΔPY1/2 deletion mutant. It is also possible that WWC1/2/3 and NF2 act synergistically to regulate YAP1 in a way that is not reflected in our experiments so far. In 293A cells, YAP1 phosphorylation in response to serum stimulation was partially inhibited by NF2 knockout and minimally inhibited by WWC1/2/3 triple knockout, but was strongly inhibited by the combined knockout of NF2 and WWC1/2/3 (59).

WWC1/2/3 may act together with NF2 to regulate YAP1 phosphorylation. Finally, our finding that in anchorage independent growth assays PTPN14 knockout promoted colony formation and reduced the levels of phosphorylation on YAP1 S127 (Figure 8) further strengthens the connection between YAP1, PTPN14, and oncogenic transformation.

Consistent with the central role of YAP1 in epithelial homeostasis and the epithelial tropic nature of HPV infection, there are additional links between HPV biology and YAP1/Hippo signaling. Cutaneous HPV8 E6 can bind TEAD and activate YAP1/TEAD-dependent transcription (101), and HPV8 E6 can also reduce LATS phosphorylation during failed cytokinesis (102). HPV E6/E7 expression in cancer cell lines downregulates MST1 by increasing miR-18A levels (103). Both high- and low-risk mucosal HPV E6 proteins have some ability to promote YAP1 nuclear localization in basal epithelial cells (64, 104). There is growing recognition that YAP1 and Hippo signaling are affected by other DNA tumor viruses. SV40 ST antigen promotes the interaction between STRIPAK and MAP4K4, thereby decreasing MAP4K4 activity and activating YAP1 (77). The Epstein-Barr Virus (EBV) protein LMP1 reduces phosphorylation on YAP1 S127, and the analogous TAZ S89 site, by increasing Src kinase activity (105).

Our findings provide the first evidence of a viral protein, HPV18 E7, targeting a Hippo pathway component to suppress LATS kinase activity, the central regulatory step immediately upstream of YAP1. PTPN14 is expressed specifically in the basal layer of stratified epithelia (64, 106), and so our finding that PTPN14 can control YAP1 and LATS phosphorylation in human keratinocytes supports that PTPN14 is a basal keratinocyte-specific regulator of YAP1 and the Hippo pathway. Our previous work supports that PTPN14 degradation and YAP1/TEAD transcriptional activity are required for the transforming activity of high-risk HPV E7 (21, 64).

Other direct interactions between HPV oncoproteins and cellular targets, such as E7/RB1 and E6/p53, are essential for HPV carcinogenesis but so far have not been tractable targets for therapeutic intervention. Understanding the mechanism by which E7 degrades PTPN14 to activate YAP1 has the potential both to provide a new therapeutic opportunity in HPV disease and to elucidate details of how HPV E7 proteins promote basal epithelial identity. Our future studies will also use E7 and PTPN14 as tools to better understand the complex interactions that control Hippo signaling and YAP1 activity in human epithelial cells.

## MATERIALS AND METHODS

### Plasmids and cloning

MSCV-puro C-terminal FlagHA retroviral expression vectors encoding HPV18 E7, HPV18 E7 R84S, or empty vector controls were previously described (21, 86). Sequences for sgRNA selected from the Broad Institute Brunello library (107) were ligated into LentiCRISPRv2 vectors following standard cloning protocols. A lentiviral vector (pHAGE) was used for PTPN14 overexpression in HFK as previously described (64). A doxycycline-inducible, HA-tagged PTPN14 overexpression vector was developed by cloning the wild-type PTPN14 coding sequence into a lentiviral vector pLIX_402, a gift from David Root (Addgene #41394), using Gateway recombination. More information for all plasmids is listed in Table S2. Mutant versions of PHAGE-PTPN14 and pLIX-PTPN14 were generated by site-directed mutagenesis and Gateway recombination.

### Cell culture

Discarded, deidentified primary human foreskin keratinocytes (HFKs) were obtained from the Skin Biology and Diseases Resource-based Center (SBDRC) at the University of Pennsylvania. N/Tert-1 keratinocytes were the gift of James Rheinwald (108). All keratinocytes were cultured as previously described (64). N/Tert-1 cell lines were established by previously described methods for lentiviral or retroviral vector expression (18, 86). Gene knockouts in N/Tert-1 cells were accomplished by one of two methods. In Figures 1 and 2, N/Tert-1 LentiCRISPR knockout cell lines were established by lentiviral transduction of LentiCRISPRv2 constructs encoding spCas9 and a sgRNA sequence selected from the Broad Institute Brunello Library for targeting PTPN14 or a nontargeting control, then selecting with G418. In Figures 4 and 6C, N/Tert-1 Cas9 knockout cells were generated by transduction with a spCas9 lentiviral expression vector (Addgene # 52962) and selecting with blasticidin, after which cells were transiently transfected with sgRNA targeting PTPN14, a non-targeting sgRNA (Synthego), or mock transfected (18).

Doxycycline inducible PTPN14 expressing N/Tert-1 keratinocytes (Figure 5) were generated by transducing N/Tert-1 cells with the lentiviral expression vector pLIX-PTPN14 and selecting with puromycin. In Figure 5, the inducible PTPN14 cells were established using parental N/Tert-1 cells. In Figure 6C, the inducible PTPN14 cells were established using N/Tert-1 Cas9:sgPTPN14 knockout cells as the starting material. N/Tert-1 Cas9:sgPTPN14 knockout cells were transduced with pLIX-PTPN14 lentiviral constructs containing the wild-type PTPN14 gene or PTPN14 ΔPY1/2, ΔPY3/4, or C1121S mutants and selected with puromycin.

Keratinocytes expressing HPV18 E7 were generated by transducing N/Tert-1 cells with MSCV-Puro retroviral vectors encoding either wild-type HPV18 E7, HPV18 E7 R84S, or empty control, then selecting with puromycin (Figure 3).

HEK TER cells were described previously and cultured in Minimum Essential Media (MEM, Gibco) supplemented with 10% fetal bovine serum (FBS), 2mM L-glutamine and penicillin-streptomycin (76). HEK TER cells expressing spCas9 were generated by transduction with a Cas9 lentiviral vector followed by blasticidin selection. HEK TER Cas9 cells were transfected with sgRNA targeting PTPN14, LATS1 and LATS2, or nontargeting control (Synthego) (18). HEK TER cells stably expressing SV40 ST were described previously (11, 76).

### Immunoblots

Immunoblots were performed using Mini-PROTEAN or Criterion (Bio-Rad) Tris/Glycine 4-20% polyacrylamide gradient SDS-PAGE gels. Proteins were transferred to polyvinylidene difluoride (PVDF) membranes as previously described (19). Membranes were blocked in 5% nonfat dried milk (NFDM) in Tris-buffered saline, pH 7.4, with 0.05% Tween 20 (TBS-T) unless otherwise specified. Blots were then incubated in 5% NFDM/TBS-T with primary antibodies listed in Table S3. Blots were incubated with anti-mouse or anti-rabbit secondary antibody conjugated to horseradish peroxidase, washed with TBS-T, and proteins detected by chemiluminescent substrate. Blots were imaged using a Chemidoc MP imaging system (BioRad). Membranes probed for YAP1 or with one of several phospho-specific antibodies (YAP1 pS127, LATS1 pT1079, NF2 pS518) were blocked with TBS-based Intercept Blocking Buffer (LI-COR) and incubated with antibodies diluted in TBS-based Intercept Antibody Diluent (LI-COR). Blots probed for LATS1 pT1079, YAP1 pS127, YAP1 and beta-actin were incubated with secondary antibody-fluorophore conjugates anti-mouse IRDye 680LT (LI-COR) or anti-rabbit IRDye 800CW (LI-COR) diluted in Intercept Antibody Diluent (LI-COR). Blots were washed with TBS-T and fluorescence was detected using a LI-COR imager. Blots probed for NF2 pS518 were incubated with anti-rabbit secondary antibody conjugated to horseradish peroxidase, diluted in Intercept Antibody Diluent (LI-COR), then washed with TBS-T and detected by chemiluminescent substrate using a Chemidoc MP imaging system (BioRad). Individual western blot bands were quantified by densitometry analysis. The mean gray pixel value of each band was determined by measuring the grey pixel value of the area containing the band of interest, then subtracting the grey pixel value from a region of the same blot that did not contain a band. The same sized area was measured for each band on a blot and for the background area on that blot. Values were measured with ImageJ software (109) (Table S1).

### Immunoprecipitation

Immunoprecipitation with anti-HA conjugated agarose beads was described previously (19). Whole cell protein lysates were incubated with anti-HA agarose beads, then washed with lysis buffer. Immunoprecipitates were eluted by boiling in protein sample buffer. Elution and input samples were analyzed by SDS-PAGE and immunoblot.

### Hippo activation assays

For cytochalasin D treatment, 2.5 x 105 cells were plated in 6 cm dishes and grown overnight before treatment with 5 µM cytochalasin D in appropriate media (MEM for HEK TER cells, K- SFM for keratinocytes). Plates were harvested every 30 minutes starting at T0 by washing once with TBS and scraping into Radioimmunoprecipitation Assay (RIPA) lysis buffer containing protease and phosphatase inhibitors. For suspension assays, 5 x 105 cells were plated in 10 cm dishes and grown for 24 hours before treatment with 0.05% Trypsin/EDTA for 10 minutes, then quenched with DMEM/F12 containing 10% FBS. Suspended cells were pelleted and washed with TBS before being pelleted again and resuspended in RIPA lysis buffer containing protease and phosphatase inhibitors. Control cells were washed once with TBS in plates and scraped into RIPA with protease and phosphatase inhibitors. All cell lysates were incubated on ice at least 30 minutes to complete lysis then protein concentrations were normalized by Bradford assay.

### Differentiation induction with calcium

N/Tert-1 mock or sgPTPN14 cells were seeded in 6 well plates at a density of 10,000 cells per well and cultured for four days. Cells were then maintained in K-SFM (untreated) or switched to culture in Keratinocyte Basal Medium (KBM, Lonza) with CaCl2 added to a final concentration of 1.5 mM. Treated and untreated cells were harvested for RNA analysis and qRT-PCR at times indicated.

### Differentiation induction with PTPN14 mutants

HFK were seeded in 12 well plates with 5.0 x 104 cells per well and transduced with pHAGE lentiviral constructs encoding GFP, wild-type PTPN14, PTPN14 ΔPY1/2, PTPN14 ΔPY3/4 or PTPN14 C1121S. Total RNA was collected 48 hours post-transduction and analyzed by qRT-PCR for KRT1 transcripts normalized to GAPDH.

### qRT-PCR

Total RNA from cell lysates was isolated with a NucleoSpin RNA purification kit (Macherey-Nagel). Purified whole cell RNA was converted to cDNA using the high-capacity cDNA reverse transcription kit (Applied Biosystems). Specific genes were quantified by qRT-PCR with Fast SYBR green master mix (Applied Biosystems) using a Quant-Studio 3 system (ThermoFisher Scientific). KiCqStart qRT-PCR primers (MilliporeSigma) were used to quantify the following transcripts: KRT1, KRT10, IVL, PTPN14, YAP1, LATS1, STK3 (MST2), STK4 (MST1), NF2, WWC1, WWC2, WWC3, and GAPDH.

### PTPN14 overexpression and siRNA knockdown assay

Primary HFK were reverse transfected with 40nM siRNAs (Dharmacon) using Dharmafect 1 transfection reagent. Transfected HFK were transduced at 24 hours post-transfection with pHAGE-PTPN14 lentivirus. At 48 hours post-transfection media was exchanged for 1:1 DF-K/K- SFM (86). Cells were lysed and total RNA collected 72 hours post-transfection for qRT-PCR analysis. The siRNAs used were: nontargeting siControl #1, siYAP1-06, siWWTR-06, siLATS1- 05, siLATS1-08, siLATS2-09, siLATS2-10, siSTK3-06, siSTK3-08, siSTK4-06, siSTK4-08, siNF2-05, siNF2-07, siWWC1-09, siWWC1-10, siWWC2-17, siWWC2-20, siWWC3-17, siWWC3-18.

### Anchorage independent growth assay

HEK TER cell lines were suspended in Dulbecco’s Modified Eagle Medium (DMEM, Gibco) and mixed with 0.4% Noble agar, then seeded onto 6 cm plates coated with DMEM/0.6% Noble agar, allowed to solidify at room temperature, then incubated at 37°C/5% CO2 (11). DMEM was supplemented with 10% FBS and antibiotic-antimycotic. Cells were seeded in triplicate plates (5.0 x 104 cells/plate) for each condition and colonies allowed to form for 18 days, then photographed with a GelDoc XR+ imager (BioRad). Colonies were counted using ImageJ software.

## ACKNOWLEDGEMENTS

We are grateful to Drs. Karl Munger and Molly Patterson for helpful comments on the manuscript and to Dr. Pushkal Ramesh for help with primer validation. We thank the members of our laboratory for helpful discussions. This work was supported by American Cancer Society grant 131661-RSG-18-048-01-MPC, NIH/NIAID R01 AI148431, and NIH/NIAID R21 AI176035 to EAW. WJB was supported by NIH/NCI T32 CA115299 and NIH/NIDCR F32 DE032573. JH was supported by NIH/NIAID T32 AI007324 and NIH/NIDCR F31 DE030365. The University of Pennsylvania SBDRC was funded by NIH/NIAMS P30 AR069589.

## AUTHOR CONTRIBUTIONS

Conception and design: WJB, JH, EAW. Acquisition of data: WJB, JH, EAW. Analysis and interpretation of data: WJB, JH, EAW. Drafting or revising the article: WJB, JH, EAW.

---

**Figure S1. PTPN14 increases KRT10 protein levels.** N/Tert-1 Cas9:sgPTPN14 keratinocytes transduced with the pLIX-PTPN14 overexpression construct were incubated with 0, 1 or 2 µg/ml doxycycline for 48 hours or 72 hours. Whole cell protein lysates were separated by SDS-PAGE and proteins were detected by immunoblotting for HA, PTPN14, KRT10, and actin.

**Figure S2. PTPN14 increases KRT1 and IVL transcript levels.** HFK transfected with siRNA for 72 hours were also transduced for 48 hours with lentiviral vectors for GFP or PTPN14 overexpression. Cells were transfected with siRNA: nontargeting control, YAP1, TAZ, and two separate pairs for LATS1 and LATS2 denoted (A) and (B). Total cellular RNA was measured by qRT-PCR for KRT1 and IVL for two biological replicate experiments, each performed in technical duplicate. Transcript levels were normalized to GAPDH. Graphs show mean ± standard deviation. Samples are the same as those analyzed in Figure 7.

**Figure S3. Validation of siRNA knockdowns.** HFK were transfected with siRNA then transduced with lentiviruses encoding GFP or PTPN14 at 24h post transfection. Total cellular RNA was collected 72h post-knockdown and 48h post-transduction. RNA transcripts for KRT10 were measured by qRT-PCR and normalized to GAPDH. Six individual experiments were conducted, each in technical duplicate, and samples are those that are analyzed in Figure 7. (A) Total cellular RNA was analyzed by qRT-PCR with primer sets as indicated to validate knockdown efficiency. Transcript levels are normalized to GAPDH. Graphs show mean RNA levels ± range. TAZ and LATS2 transcript levels were below the limit of detection and are not shown. (B) PTPN14 transcripts were measured in all six experiments by qRT-PCR, normalized to GAPDH, and plotted as mean ± range.

**Figure S4. PTPN14 knockout promotes anchorage independent growth and suppresses YAP1 phosphorylation in HEK TER cells.** HEK TER cells expressing Cas9 were transfected with a second set of sgRNA compared to those used in Figure 8. HEK TER cells expressing SV40 ST were used as a positive control for colony formation. Cells were plated in soft agar in technical triplicates and incubated at 37°C for 18 days, then photographed. Colonies were counted and quantified using ImageJ software. Graphs show individual data points for each plate and indicate mean ± standard deviation. Statistical significance of the nontargeting control condition compared to experimental conditions was determined by ANOVA with Dunnett’s multiple comparisons test (ns, not significant; ****, P < 0.0001).

Supplementary Table 1. Densitometry analysis of western blots in Figures 1, 2, 3, and 8.

Supplementary Table 2. List of plasmids used in the study.

Supplementary Table 3. List of antibodies used in the study.

