## Supplemental Table 1 for "HPV18 E7 inhibits LATS1 kinase and activates YAP1 by degrading PTPN14"

**Figure 1**

| | sgNT-1 | | | | | | | |
|---|---|---|---|---|---|---|---|---|
| CytoD (min): | 0 | backgroun | 30 | backgroun | 60 | backgroun | 90 | backgroun |
| pYAP | 95.303 | 44.068 | 134.949 | 49.318 | 129.412 | 49.333 | 149.736 | 60.071 |
| YAP | 65.683 | 30.969 | 74.743 | 34.251 | 66.27 | 30.603 | 66.386 | 30.608 |
| pYAP | 92.578 | 33.268 | 105.273 | 36.367 | 112.567 | 36.308 | 113.064 | 36.671 |
| YAP | 83.408 | 55.021 | 82.099 | 47.664 | 71.585 | 37.286 | 71.832 | 40.654 |
| pYAP | 110.828 | 53.521 | 133.436 | 58.713 | 133.952 | 53.404 | 160.787 | 56.62 |
| YAP | 143.04 | 117.467 | 130.703 | 83.014 | 123.591 | 71.905 | 124.772 | 80.489 |

| | sgNT-2 | | | | | | | |
|---|---|---|---|---|---|---|---|---|
| CytoD (min): | 0 | backgroun | 30 | backgroun | 60 | backgroun | 90 | backgroun |
| pYAP | 98.969 | 50.187 | 117.255 | 53.26 | 116.279 | 53.82 | 141.542 | 53.976 |
| YAP | 65.524 | 32.752 | 66.395 | 30.681 | 64.994 | 35.158 | 65.891 | 32.897 |
| pYAP | 96.686 | 35.882 | 94.666 | 40.016 | 103.062 | 38.516 | 128.11 | 41.205 |
| YAP | 73.554 | 40.901 | 75.527 | 44.527 | 73.508 | 40.581 | 77.888 | 40.478 |
| pYAP | 134.693 | 56.284 | 154.433 | 60.031 | 143.515 | 57.859 | 152.316 | 58.997 |
| YAP | 138.742 | 96.147 | 145.314 | 104.091 | 135.373 | 91.89 | 129.71 | 92.697 |

| | sgNT-1 | | | | sgPTPN14-1 | | | |
|---|---|---|---|---|---|---|---|---|
| CytoD (min): | 0 | 30 | 60 | 90 | 0 | 30 | 60 | 90 |
| pYAP/YAP | 1.475917 | 2.114763 | 2.245185 | 2.506149 | 1.375056 | 1.540785 | 1.859305 | 1.817944 |
| pYAP/YAP | 2.089337 | 2.001045 | 2.223359 | 2.450221 | 1.146721 | 1.04133 | 1.001112 | 1.240461 |
| pYAP/YAP | 2.240918 | 1.566881 | 1.55841 | 2.352302 | 1.020363 | 1.533598 | 1.769807 | 1.981521 |

**Figure 2A**

| | Untreated | | | | Suspension | | | |
|---|---|---|---|---|---|---|---|---|
| | sgNT-1 | backgroun | sgPTPN14- | backgroun | sgNT-1 | backgroun | sgPTPN14- | backgroun |
| pYAP | 68.777 | 36.141 | 65.497 | 36.858 | 129.537 | 48.027 | 99.97 | 39.908 |
| YAP | 102.93 | 76.023 | 98.933 | 72.108 | 99.841 | 71.113 | 102.535 | 73.171 |
| pLATS1 | 117.085 | 105.495 | 119.055 | 115.555 | 172.38 | 123.538 | 161.298 | 147.89 |
| LATS1 | 177.479 | 130.268 | 177.602 | 129.867 | 181.014 | 130.426 | 180.368 | 130.209 |
| pNF2 | 80.429 | 26.493 | 86.925 | 27.636 | 57.142 | 28.366 | 57.151 | 25.617 |
| NF2 | 78.661 | 18.603 | 72.345 | 11.8 | 89.663 | 11.867 | 87.679 | 15.401 |
| pYAP | 48.776 | 34.169 | 44.075 | 34.448 | 209.361 | 70.576 | 60.789 | 38.491 |
| YAP | 88.585 | 68.323 | 83.202 | 66.928 | 112.486 | 76.188 | 77.182 | 67.471 |
| pLATS1 | 134.411 | 134.371 | 134.864 | 133.557 | 233.918 | 166.639 | 193.77 | 183.397 |
| LATS1 | 162.84 | 129.07 | 155.812 | 127.962 | 206.061 | 134.651 | 154.185 | 127.467 |
| pNF2 | 207.226 | 177.823 | 201.37 | 179.366 | 206.513 | 183.535 | 182.262 | 174.584 |
| NF2 | 181.947 | 115.065 | 162.294 | 108.574 | 195.134 | 117.7 | 172.227 | 103.363 |
| pYAP | 75.971 | 35.997 | 51.796 | 31.42 | 177.681 | 42.938 | 90.313 | 40.183 |
| YAP | 77.429 | 44.703 | 65.529 | 44.282 | 91.283 | 49.26 | 69.505 | 47.77 |
| pLATS1 | 122.779 | 119.771 | 129.302 | 124.946 | 183.826 | 140.474 | 144.712 | 129.446 |
| LATS1 | 123.064 | 52.671 | 91.275 | 54.371 | 145.509 | 65.703 | 106.192 | 55.044 |
| pNF2 | 169.283 | 82.455 | 145.24 | 75.146 | 132.314 | 76.225 | 102.362 | 74.883 |
| NF2 | 119.222 | 83.063 | 128.167 | 80.773 | 171.173 | 85.8 | 149.321 | 81.707 |

**Figure 2B**

| | Untreated | | | | Suspension | | | |
|---|---|---|---|---|---|---|---|---|
| | sgNT-2 | backgroun | sgPTPN14- | backgroun | sgNT-2 | backgroun | sgPTPN14- | backgroun |
| pYAP | 131.684 | 83.432 | 107.042 | 70.532 | 186.111 | 68.577 | 135.365 | 74.66 |
| YAP | 104.428 | 73.547 | 92.393 | 68.102 | 105.909 | 67.644 | 102.016 | 67.198 |
| pLATS1 | 126.382 | 113.68 | 114.615 | 108.617 | 143.861 | 108.501 | 118.14 | 109.903 |
| LATS1 | 201.244 | 136.797 | 197.171 | 134.16 | 207.006 | 136.135 | 198.907 | 137.947 |
| pNF2 | 148.377 | 77.516 | 138.814 | 82.179 | 117.564 | 86.121 | 117.581 | 89.61 |
| NF2 | 155.092 | 81.208 | 152.42 | 77.268 | 164.197 | 78.586 | 162.285 | 73.538 |
| pYAP | 133.808 | 53.184 | 113.607 | 46.548 | 183.054 | 57.795 | 140.797 | 55.937 |
| YAP | 90.483 | 55.763 | 88.73 | 52.381 | 80.199 | 49.266 | 89.905 | 56.373 |
| pLATS1 | 100.996 | 90.073 | 101.114 | 96.722 | 139.553 | 102.692 | 121.551 | 103.571 |
| LATS1 | 198.289 | 118.126 | 194.281 | 114.444 | 191.537 | 115.274 | 195.622 | 113.453 |
| pNF2 | 101.861 | 43.413 | 108.409 | 48.409 | 76.525 | 45.683 | 75.262 | 42.641 |
| NF2 | 87.821 | 21.927 | 79.863 | 24.584 | 86.839 | 25.463 | 93.321 | 25.744 |
| pYAP | 75.306 | 37.444 | 48.099 | 32.911 | 147.176 | 46.011 | 105.255 | 36.057 |
| YAP | 73.508 | 45.927 | 64.675 | 44.526 | 72.335 | 43.658 | 78.554 | 46.232 |
| pLATS1 | 135.985 | 127.792 | 116.306 | 114.322 | 171.838 | 112.807 | 137.974 | 120.685 |
| LATS1 | 122.187 | 57.354 | 98.016 | 52.697 | 135.16 | 55.751 | 131.619 | 61.279 |
| pNF2 | 161.224 | 82.187 | 147.608 | 75.781 | 106.046 | 73.158 | 112.563 | 75.783 |
| NF2 | 145.801 | 88.374 | 137.892 | 81.288 | 158.71 | 79.408 | 163.599 | 80.133 |

| | Figure 2A | | | | Figure 2B | | | |
|---|---|---|---|---|---|---|---|---|
| | Untreated | | Suspension | | Untreated | | Suspension | |
| | sgNT-1 | sgPTPN14- | sgNT-1 | sgPTPN14- | sgNT-2 | sgPTPN14- | sgNT-2 | sgPTPN14- |
| pYAP/YAP | 1.212919 | 1.067623 | 2.837302 | 2.04543 | 1.562514 | 1.503026 | 3.07158 | 1.743495 |
| pLATS1/LATS1 | 0.245494 | 0.073321 | 0.965486 | 0.26731 | 0.197092 | 0.09519 | 0.498935 | 0.135121 |
| pNF2/NF2 | 0.898065 | 0.979255 | 0.36989 | 0.436288 | 0.959085 | 0.753606 | 0.367278 | 0.315177 |
| pYAP/YAP | 0.720906 | 0.591557 | 3.823489 | 2.296159 | 2.32212 | 1.844865 | 4.049365 | 2.530717 |
| pLATS1/LATS1 | 0.001184 | 0.04693 | 0.942151 | 0.38824 | 0.13626 | 0.055012 | 0.483341 | 0.218817 |
| pNF2/NF2 | 0.439625 | 0.409605 | 0.296743 | 0.111495 | 0.887 | 1.085403 | 0.502509 | 0.482723 |
| pYAP/YAP | 1.221475 | 0.959006 | 3.206411 | 2.306418 | 1.372757 | 0.753784 | 3.52774 | 2.140895 |
| pLATS1/LATS1 | 0.042732 | 0.118036 | 0.543217 | 0.298467 | 0.126371 | 0.043779 | 0.743379 | 0.245792 |
| pNF2/NF2 | 2.401283 | 1.478964 | 0.656988 | 0.40641 | 1.376304 | 1.268939 | 0.414718 | 0.440658 |

| | Empty Vector | | | | | | | |
|---|---|---|---|---|---|---|---|---|
| CytoD (min): | 0 | backgroun | 30 | backgroun | 60 | backgroun | 90 | backgroun |
| pYAP | 95.481 | 36.311 | 118.148 | 35.408 | 178.166 | 44.2 | 185.098 | 40.987 |
| YAP | 79.159 | 35.539 | 72.925 | 31.477 | 87.165 | 36.819 | 82.064 | 32.971 |
| pYAP | 115.789 | 39.913 | 107.597 | 39.746 | 97.508 | 35.667 | 108.077 | 38.324 |
| YAP | 53.781 | 30.465 | 55.299 | 31.592 | 48.995 | 30.091 | 45.427 | 30.249 |
| pYAP | 120.326 | 36.68 | 149.19 | 45.818 | 172.591 | 49.611 | 193.235 | 59.809 |
| YAP | 82.946 | 63.677 | 85.664 | 66.419 | 85.375 | 66.231 | 86.72 | 68.064 |
| | EmptyV | | | | 18E7 | | | |
| CytoD (min): | 0 | 30 | 60 | 90 | 0 | 30 | 60 | 90 |
| pYAP/YAP | 1.356488 | 1.996236 | 2.660907 | 2.935469 | 1.390787 | 1.439709 | 1.168746 | 1.276229 |
| pYAP/YAP | 3.254246 | 2.862066 | 3.271318 | 4.595665 | 1.749755 | 1.427272 | 0.820016 | 1.4532 |

| | | | | | | | | |
|---|---|---|---|---|---|---|---|---|
| pYAP/YAP | 4.340962 | 5.371369 | 6.423945 | 7.151908 | 3.426362 | 2.290689 | 1.95338 | 2.536999 |

**Figure 3B**

| | Untreated | | | | | | | |
|---|---|---|---|---|---|---|---|---|
| | Empty | backgroun | 18E7 | backgroun | 18E7 R84S | backgroun | Empty | backgroun |
| pYAP | 88.758 | 31.213 | 62.958 | 31.908 | 77.066 | 31.994 | 171.168 | 38.742 |
| YAP | 71.965 | 45.84 | 71.822 | 41.189 | 68.447 | 40.266 | 72.27 | 33.117 |
| pLATS1 | 43.577 | 40.719 | 40.943 | 57.649 | 50.378 | 67.831 | 123.273 | 71.652 |
| LATS1 | 127.565 | 73.201 | 129.633 | 73.748 | 131.786 | 71.384 | 131.559 | 71.03 |
| pYAP | 121.936 | 48.81 | 177.542 | 59.118 | 189.315 | 61.773 | 230.115 | 72.566 |
| YAP | 157.376 | 115.482 | 183.375 | 119.619 | 180.261 | 121.01 | 175.731 | 119.438 |
| pLATS1 | 127.597 | 124.749 | 133.062 | 141.062 | 139.366 | 129.443 | 177.971 | 138.619 |
| LATS1 | 103.572 | 43.675 | 138.787 | 46.139 | 125.722 | 44.626 | 131.561 | 42.345 |
| pYAP | 137.035 | 58.611 | 97.15 | 56.072 | 134.657 | 59.447 | 202.275 | 65.401 |
| YAP | 202.143 | 136.378 | 184.162 | 135.842 | 186.944 | 131.027 | 185.666 | 132.896 |
| pLATS1 | 110.387 | 104.633 | 107.085 | 102.99 | 112.995 | 106.235 | 156.259 | 114.724 |
| LATS1 | 180.313 | 87.789 | 165.151 | 88.852 | 163.972 | 85.367 | 163.032 | 90.104 |

| | Untreated | | | Suspension | | |
|---|---|---|---|---|---|---|
| | EmptyV | 18E7 | 18E7 R84S | EmptyV | 18E7 | 18E7 R84S |
| pYAP/YAP | 2.202679 | 1.013613 | 1.599375 | 3.38227 | 1.75801 | 3.145487 |
| pLATS1/LATS1 | 0.052572 | -0.29894 | -0.28895 | 0.852831 | 0.206883 | 0.922873 |
| pYAP/YAP | 1.745501 | 1.857457 | 2.152571 | 2.798732 | 1.760412 | 3.321504 |
| pLATS1/LATS1 | 0.047548 | -0.08635 | 0.122361 | 0.441087 | 0.110195 | 0.561575 |
| pYAP/YAP | 1.192488 | 0.850124 | 1.345029 | 2.593784 | 1.316299 | 2.981392 |
| pLATS1/LATS1 | 0.062189 | 0.05367 | 0.086 | 0.569534 | 0.17562 | 0.513546 |

**Figure 8**

| | sgNT-1 | | | | | | | |
|---|---|---|---|---|---|---|---|---|
| | 0 | backgroun | 30 | backgroun | 60 | backgroun | 90 | backgroun |
| pYAP | 113.227 | 83.682 | 147.77 | 92.754 | 145.061 | 91.637 | 158.317 | 96.746 |
| YAP | 128.566 | 96.925 | 119.956 | 81.758 | 102.337 | 70.51 | 108.233 | 64.623 |
| pYAP | 127.559 | 96.142 | 151.516 | 99.928 | 160.08 | 93.455 | 164.591 | 91.648 |
| YAP | 161.569 | 121.889 | 157.559 | 129.039 | 147.884 | 123.055 | 144.112 | 120.825 |
| | sgLATS1/2-1 | | | | | | | |
| | 0 | backgroun | 30 | backgroun | 60 | backgroun | 90 | backgroun |
| pYAP | 102.003 | 81.804 | 131.921 | 90.272 | 133.38 | 90.299 | 132.855 | 89.913 |
| YAP | 99.885 | 59.732 | 111.764 | 61.122 | 101.887 | 54.021 | 97.292 | 56.711 |
| pYAP | 115.236 | 90.996 | 149.673 | 92.467 | 147.802 | 93.137 | 150.269 | 95.105 |
| YAP | 159.816 | 128.699 | 163.897 | 123.527 | 160.555 | 128.842 | 162.213 | 123.339 |

| | sgNT-1 | | | | sgPTPN14-1 | | | |
|---|---|---|---|---|---|---|---|---|
| | 0 | 30 | 60 | 90 | 0 | 30 | 60 | 90 |
| pYAP/YAP | 0.933757 | 1.440285 | 1.678575 | 1.411855 | 0.631365 | 0.851854 | 0.973989 | 1.07125 |
| pYAP/YAP | 0.791759 | 1.808836 | 2.683354 | 3.132349 | 1.064157 | 1.460521 | 1.516648 | 1.615688 |

| sgPTPN14-1 | | | | | | | |
|---|---|---|---|---|---|---|---|
| 0 | backgroun | 30 | backgroun | 60 | backgroun | 90 | background |
| 92.538 | 46.486 | 103.181 | 48.252 | 119.552 | 49.168 | 116.625 | 56.162 |
| 63.634 | 30.143 | 67.683 | 32.033 | 69.075 | 31.22 | 63.74 | 30.481 |
| 63.536 | 31.867 | 70.302 | 32.836 | 63.363 | 31.862 | 69.715 | 34.115 |
| 64.216 | 36.599 | 71.834 | 35.855 | 65.764 | 34.298 | 66.337 | 37.638 |
| 93.563 | 50.77 | 127.824 | 54.334 | 149.492 | 55.894 | 143.859 | 55.713 |
| 129.187 | 87.248 | 137.757 | 89.837 | 145.141 | 92.255 | 143.328 | 98.844 |

| sgPTPN14-3 | | | | | | | |
|---|---|---|---|---|---|---|---|
| 0 | backgroun | 30 | backgroun | 60 | backgroun | 90 | background |
| 80.441 | 42.689 | 99.469 | 44.297 | 93.607 | 43.249 | 101.658 | 45.206 |
| 64.036 | 34.181 | 71.187 | 31.987 | 67.211 | 35.057 | 62.911 | 34.736 |
| 83.453 | 34.527 | 68.569 | 32.617 | 67.516 | 34.553 | 59.507 | 34.824 |
| 71.569 | 36.997 | 67.287 | 32.897 | 71.08 | 42.083 | 66.39 | 41.411 |
| 108.125 | 53.651 | 108.735 | 54.194 | 123.621 | 55.727 | 101.688 | 51.069 |
| 128.311 | 91.164 | 129.368 | 92.057 | 136.012 | 91.042 | 135.131 | 109.094 |

| sgNT-2 | | | | sgPTPN14-3 | | | |
|---|---|---|---|---|---|---|---|
| 0 | 30 | 60 | 90 | 0 | 30 | 60 | 90 |
| 1.488527 | 1.791874 | 2.093411 | 2.653998 | 1.264512 | 1.407449 | 1.56615 | 2.00362 |
| 1.862126 | 1.762903 | 1.960276 | 2.323042 | 1.415191 | 1.04542 | 1.136773 | 0.98815 |
| 1.840803 | 2.290032 | 1.969873 | 2.521249 | 1.466444 | 1.461794 | 1.509762 | 1.944118 |

d

d

3

**Figure 3A**

| 18E7 | | | | | | | | | |
|---|---|---|---|---|---|---|---|---|---|
| 0 | backgroun | 30 | backgroun | 60 | backgroun | 90 | backgroun | 0 | backgroun |
| 131.684 | 37.97 | 112.693 | 34.786 | 88.179 | 32.826 | 101.809 | 35.283 | 116.162 | 37.281 |
| 109.109 | 41.727 | 85.561 | 31.448 | 81.245 | 33.884 | 86.335 | 34.208 | 79.509 | 35.254 |
| 66.897 | 32.978 | 56.254 | 31.458 | 52.367 | 34.284 | 75.271 | 36.302 | 91.747 | 37.468 |
| 49.674 | 30.289 | 47.592 | 30.219 | 52.839 | 30.787 | 57.152 | 30.336 | 56.919 | 32.933 |
| 103.65 | 40.742 | 93.09 | 42.239 | 74.436 | 37.564 | 95.035 | 40.591 | 111.466 | 42.384 |
| 86.86 | 68.5 | 93.048 | 70.849 | 90.019 | 71.143 | 91.453 | 69.993 | 91.697 | 71.246 |

| 18E7 R84S | | | |
|---|---|---|---|
| 0 | 30 | 60 | 90 |
| 1.78242 | 2.239326 | 2.768147 | 2.92193 |
| 2.262945 | 1.525043 | 2.087527 | 3.159851 |

| | | | |
|---|---|---|---|
| 3.377928 | 2.497191 | 2.924334 | 4.150392 |

| Suspension | | | |
|---|---|---|---|
| 18E7 | backgroun | 18E7 R84S | background |
| 94.55 | 32.276 | 151.452 | 37.296 |
| 65.443 | 30.02 | 71.238 | 34.946 |
| 74.118 | 61.073 | 120.633 | 61.056 |
| 133.197 | 70.142 | 136.463 | 71.907 |
| 157.94 | 57.211 | 211.29 | 64.443 |
| 177.256 | 120.037 | 163.122 | 118.911 |
| 149.304 | 139.959 | 177.085 | 140.285 |
| 128.6 | 43.796 | 109.716 | 44.186 |
| 113.209 | 57.365 | 193.126 | 64.151 |
| 180.882 | 138.457 | 186.413 | 143.153 |
| 126.602 | 114.235 | 155.027 | 114.064 |
| 160.551 | 90.132 | 166.78 | 87.015 |

| sgPTPN14-1 | | | | | | | |
|---|---|---|---|---|---|---|---|
| 0 | backgroun | 30 | backgroun | 60 | backgroun | 90 | background |
| 108.148 | 82.418 | 127.76 | 86.802 | 146.453 | 93.13 | 142.778 | 94.726 |
| 108.954 | 68.201 | 117.535 | 69.454 | 120.997 | 66.25 | 107.366 | 62.51 |
| 117.822 | 90.454 | 138.946 | 91.926 | 144.967 | 91.399 | 152.264 | 94.571 |
| 149.88 | 124.162 | 152.072 | 119.878 | 160.661 | 125.341 | 158.093 | 122.385 |
| SV40ST | | | | | | | |
| 0 | backgroun | 30 | backgroun | 60 | backgroun | 90 | background |
| 89.902 | 78.266 | 115.911 | 84.392 | 128.08 | 87.836 | 121.006 | 83.218 |
| 93.412 | 50.293 | 80.315 | 49.646 | 104.592 | 47.388 | 105.051 | 75.542 |
| 106.939 | 89.705 | 133.043 | 90.5 | 134.944 | 93.265 | 129.483 | 91.79 |
| 150.837 | 121.157 | 159.717 | 121.805 | 154.162 | 119.693 | 166.791 | 131.333 |

| sgLATS1/2-1 | | | | SV40 ST | | | |
|---|---|---|---|---|---|---|---|
| 0 | 30 | 60 | 90 | 0 | 30 | 60 | 90 |
| 0.503051 | 0.82242 | 0.900033 | 1.05818 | 0.269858 | 1.027715 | 0.703517 | 1.280558 |
| 0.778995 | 1.417042 | 1.723741 | 1.419046 | 0.58066 | 1.122151 | 1.209173 | 1.063032 |

| 18E7 R84S | | | | | |
|---|---|---|---|---|---|
| 30 | backgroun | 60 | backgroun | 90 | background |
| 138.238 | 40.001 | 127.705 | 38.698 | 150.022 | 40.698 |
| 78.477 | 34.608 | 69.032 | 36.878 | 70.915 | 33.5 |
| 76.636 | 39.123 | 79.901 | 36.303 | 96.187 | 36.766 |
| 57.098 | 32.5 | 57.542 | 36.657 | 56.529 | 37.724 |
| 101.056 | 42.372 | 104.273 | 41.084 | 146.479 | 43.79 |
| 90.729 | 67.229 | 87.128 | 65.52 | 87.975 | 63.233 |
