## Supplemental Table 2 for "HPV18 E7 inhibits LATS1 kinase and activates YAP1 by degrading PTPN14"

| Name | Lab ID # | Addgene # | Gene | Promoter | Bacterial Resistance | Selection | Epitope Tag | Tag Location | Source |
|---|---|---|---|---|---|---|---|---|---|
| MSCV-IP N HA only empty v2 | 7270 | 163302 | N/A | MSCV LTR | Ampicillin | Puromycin | HA | N-terminus | White et al (2014) J. Virol. 88(15):8201-12 |
| MSCV-P C-FlagHA 18E7 | 6641 | 35019 | HPV18 E7 | MSCV LTR | Ampicillin | Puromycin | FlagHA | C-terminus | White et al. (2012) PNAS: 109(5):E260–E267 |
| MSCV-P C-FlagHA 18E7 R84S | 8193 | 163307 | HPV18 E7 R84S | MSCV LTR | Ampicillin | Puromycin | FlagHA | C-terminus | Hatterschide et al. (2020) J. Virol. 94:e1024-20 |
| pLIX-402 | 8201 | 41394 | N/A | TRE | Ampicillin+chloramphenicol | N/A | HA | C-terminus | Addgene |
| pLIX-PTPN14 | 8221 | 221643 | PTPN14 | TRE | Ampicillin | Puromycin | HA | C-terminus | This study |
| pLIX-PTPN14 ΔPPXY1/2 | 8224 | 221644 | PTPN14 | TRE | Ampicillin | Puromycin | HA | C-terminus | This study |
| pLIX-PTPN14 ΔPPXY3/4 | 8552 | 221645 | PTPN14 | TRE | Ampicillin | Puromycin | HA | C-terminus | This study |
| LentiCas9-blast | 7527 | 52962 | spCas9 | EFS-NS | Ampicillin | Blasticidin | | N-terminus | Addgene |
| pHAGE-P-CMVt N-HA GFP | 6571 | N/A | GFP | CMVt | Ampicillin | Puromycin | HA | N-terminus | Galligan et al. (2015) J Proteome Res. 14(2): 953–966. |
| pHAGE-P-CMVt N-V5 PTPN14 | 7522 | N/A | PTPN14 | CMVt | Ampicillin | Puromycin | V5 | N-terminus | White et al. (2016) mBio. 7(5):e01530-16 |
| pHAGE-P-CMVt N-V5 PTPN14 ΔPPXY1/2 | 8215 | 221646 | PTPN14 | CMVt | Ampicillin | Puromycin | V5 | N-terminus | This study |
| pHAGE-P-CMVt N-V5 PTPN14 ΔPPXY3/4 | 8180 | 221647 | PTPN14 | CMVt | Ampicillin | Puromycin | V5 | N-terminus | This study |
| pHAGE-P-CMVt N-V5 PTPN14 C1121S | 8189 | 221648 | PTPN14 | CMVt | Ampicillin | Puromycin | V5 | N-terminus | This study |

| Name | Lab ID # | Addgene # | sgRNA sequence | Promoter | Bacterial Resistance | Selection | Source |
|---|---|---|---|---|---|---|---|
| LentiCRISPRv2-Neo | 8175 | 98292 | N/A | U6 | Ampicillin | G418 | Addgene |
| LentiCRISPRv2 Neo sgNT-1 | 8389 | 221649 | GAGCTCGCCATG | U6 | Ampicillin | G418 | This study |
| LentiCRISPRv2 Neo sgNT-2 | 8390 | 221650 | GGATTGTGGTCG | U6 | Ampicillin | G418 | This study |
| LentiCRISPRv2 Neo sgPTPN14-1 | 8391 | 221651 | CATGACTGTCTC | U6 | Ampicillin | G418 | This study |
| LentiCRISPRv2 Neo sgPTPN14-3 | 8392 | 221652 | CCACACTGGACG | U6 | Ampicillin | G418 | This study |
