## Supplemental Table 3 for "HPV18 E7 inhibits LATS1 kinase and activates YAP1 by degrading PTPN14"

| Target | Description | Company | Catalog # | Application | Dilution |
|---|---|---|---|---|---|
| Beta-Actin | Mouse anti-bActin | CST | 3700S | Western Blot | 1:1000 |
| PTPN14 | Rabbit anti-PTPN14 | CST | 13808S | Western Blot | 1:1000 |
| YAP1 | Mouse anti-YAP1 | CST | 12395S | Western Blot | 1:1000 |
| YAP1 pS127 | Rabbit anti-YAP1 pS127 | CST | 4911S | Western Blot | 1:1000 |
| LATS1 | Mouse anti-LATS1 | Proteintech | 66569-1-IG | Western Blot | 1:1000 |
| LATS1 pT1079 | Rabbit anti-LATS1 pT1079 | CST | 8654S | Western Blot | 1:1000 |
| NF2/Merlin | Rabbit anti-Merlin | CST | 12888S | Western Blot | 1:1000 |
| NF2 pS518 | Rabbit anti-Merlin pS518 | CST | 13281 | Western Blot | 1:1000 |
| RB1 | Mouse anti-RB1 | CalBiochem | OP66 | Western Blot | 1:500 |
| HA | Rat anti-HA HRP conjugate | Roche | 12013819001 | Western Blot | 1:500 |
| WWC1/Kibra | Rabbit anti-Kibra | CST | 8774S | Western Blot | 1:1000 |
| Keratin 10 | Mouse anti Cytokeratin 10 | Santa Cruz | sc-52318 | Western Blot | 1:1000 |
| Mouse IgG | Horse anti-Mouse HRP conjugate | CST | 7076S | Western Blot | 1:2000 |
| Rabbit IgG | Horse anti-Rabbit HRP conjugate | CST | 7074S | Western Blot | 1:2000 |
| Mouse IgG | Goat anti-Mouse IRDye 680LT | LI-COR | 926-68020 | Western Blot | 1:5000 |
| Rabbit IgG | Goat anti-Rabbit IRDye 800CW | LI-COR | 926-32211 | Western Blot | 1:5000 |
