## Supplementary figures and images for "HPV18 E7 inhibits LATS1 kinase and activates YAP1 by degrading PTPN14"

### Supplemental Figure 1

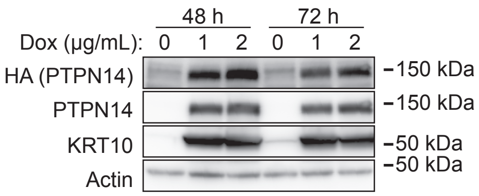

### Supplemental Table 2

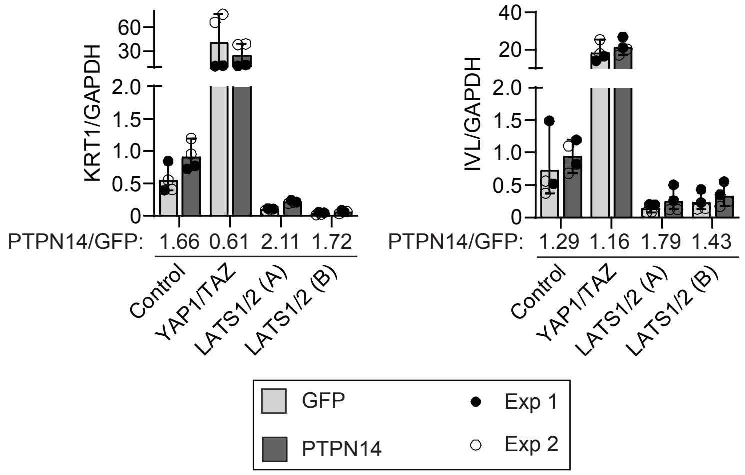

### Supplemental Table 3

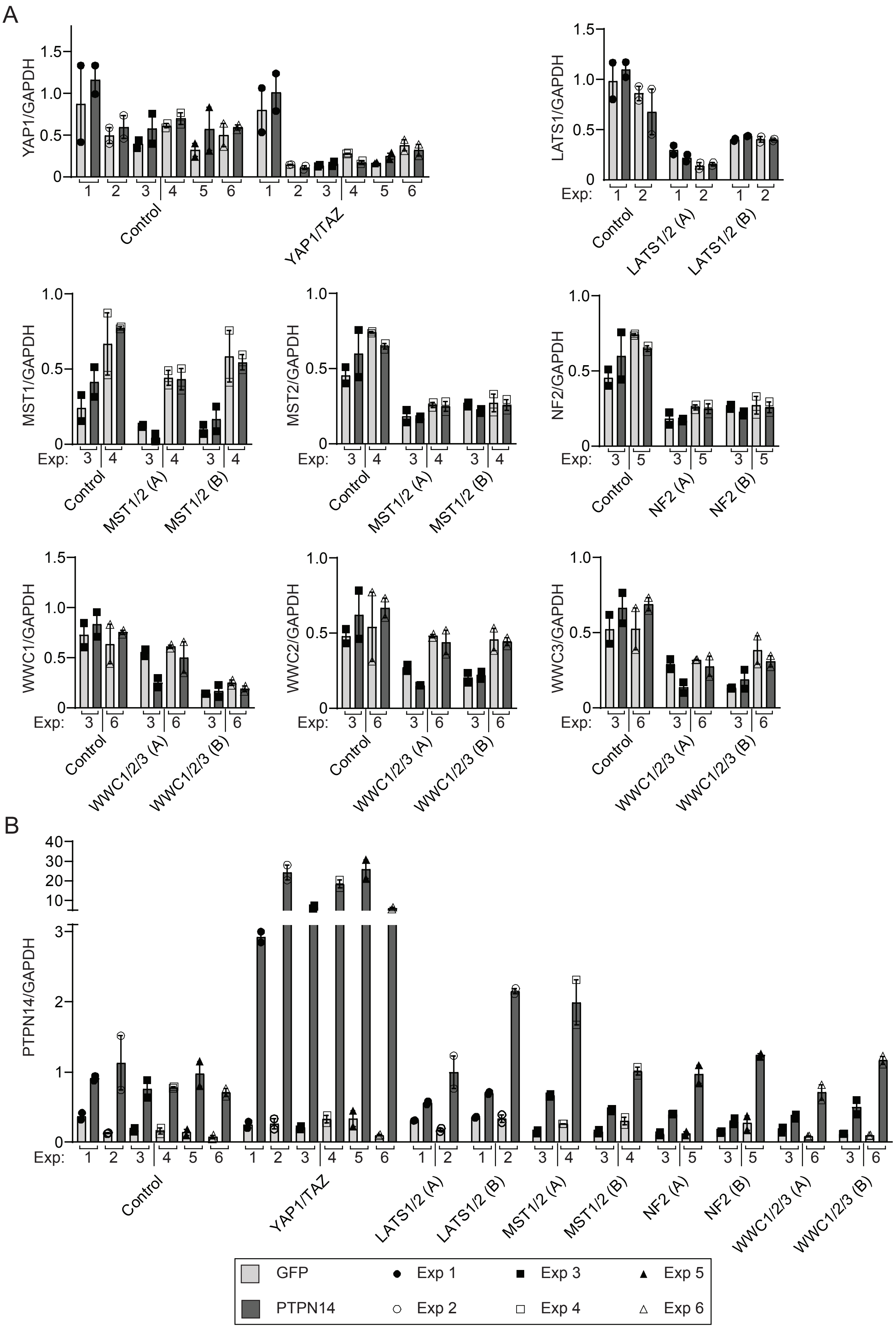

### Supplemental Table 4

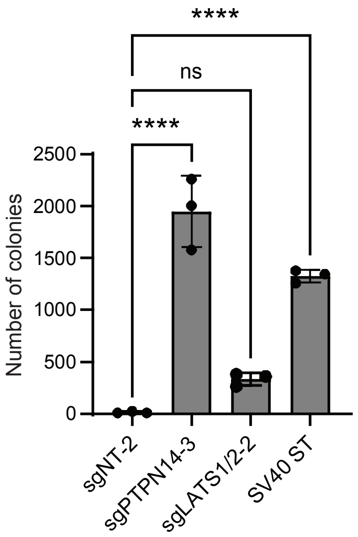
